# Follicle Stem Cells (FSCs) in the Drosophila ovary; a critique of published studies defining the number, location and behavior of FSCs

**DOI:** 10.1101/2020.06.25.171579

**Authors:** Daniel Kalderon, David Melamed, Amy Reilein

## Abstract

A paper by Reilein et al., (2017) presented several key new insights into the behavior of adult Follicle Stem Cells (FSCs) in the Drosophila ovary, including overwhelming evidence that each ovariole hosts a large number of FSCs (about 14-16) maintained by population asymmetry (Reilein et al., 2017), rather than just two FSCs, dividing with largely individually asymmetric outcomes, as originally proposed (Margolis and Spradling, 1995; Nystul and Spradling, 2007). Here we provide further discussion asserting the merits of the conclusions of Reilein et al., (2017) and the deficiencies in the contrary assertions recently presented by Fadiga and Nystul (Fadiga and Nystul, 2019). The principles that we discuss here, particularly with regard to lineage tracing and population asymmetry, are common to the investigation of most types of adult stem cell and should therefore be instructive and of interest to investigators studying any type of adult stem cell. The improved understanding of FSC numbers, location and behavior afforded by Reilein et al., (2017) and Reilein et al., (2018) can only provide a firm foundation for future progress once they are widely appreciated and seen to be resistant to challenge, as described in detail here.

## Introduction

In 2017 our group published a paper entitled, “Alternative direct stem cell derivatives defined by stem cell location and graded Wnt signaling” (Reilein et al., 2017). The paper included at least five major discoveries. These included two major revisions of our understanding of the identity and behavior of Drosophila ovarian Follicle Stem Cells (FSCs); (a) the existence of 14-16, rather than two or three FSCs, and (b) that these FSCs, which exist in all radial locations within a narrow anterior-posterior (AP) domain of each germarium, are maintained by population asymmetry with frequent loss and amplification of individual lineages rather than repeated asymmetric division outcomes for each FSC. The other major discoveries were that (a) FSCs, defined by the production of proliferative Follicle Cells (FCs), also produce non-proliferative Escort Cells (ECs), which support early germline differentiation in the anterior half of the germarium, (b) posteriorly located FSCs directly produce (or more accurately, as shown in (Reilein et al., 2018), become) FCs while anterior FSCs produce ECs but FSCs also change AP locations relatively frequently so that a single FSC lineage can produce both FCs and ECs, and (c) the strength of Wnt signaling, which declines from anterior to posterior across the FSC domain, strongly influences FSC location and the propensity to produce ECs or FCs. The latter three discoveries were emphasized in the title, summary and discussion in order to draw the attention to insights that were new, and therefore of most relevance, for the whole broad field of adult stem cell biology.

Fadiga and Nystul now challenge the first two, fundamental discoveries of Reilein et al., (2017) concerning FSC numbers, locations and organization, proposing instead that the very first model of two relatively long-lived FSCs, repeatedly undergoing divisions with asymmetric outcomes is correct (Margolis and Spradling, 1995; Fadiga and Nystul, 2019). Despite the rhetoric alleging problems with the conclusions of Reilein et al., (2017), Fadiga and Nystul do not present a single valid criticism concerning any of the conclusions, large or small, in Reilein et al., (2017) and they present no new lines of investigation or insights. First, Fadiga and Nystul present a multicolor lineage analysis using genetic reagents that we made and used in the studies of Reilein et al., (2017) and then supplied to Dr. Nystul. Fadiga and Nystul present flawed experiments and analysis using these reagents. Rather than seeking to rectify these flaws, some of which they specifically outline, Fadiga and Nystul claim the results support a lower number of FSCs and that the genetic system is unreliable; hence, they argue that the conclusions of Reilein et al., (2017) should be doubted. However, the analysis of Reilein et al., (2017) did not have any of the experimental or logical flaws of the multicolor lineage analysis of Fadiga and Nystul that led them to incorrect conclusions. The genetic reagents and reasoning of Reilein et al., (2017) were entirely sound and every conclusion from that multicolor clonal analysis remains valid.

Fadiga and Nystul then present single-color lineage analyses of the type that has been performed hundreds of times in our laboratory and other laboratories over the last twenty years. They report that an FSC lineage almost always contributes FCs to each new egg chamber, that a single FSC contributes about half of the FCs in an ovariole and that the FSCs are located within a single AP plane and at only a fraction of all radial locations. However, these conclusions are based on misrepresentation of their own data, on a circular argument that selects for scoring only clones that conform to a preferred outcome, and on imaging that does not have the clarity to reveal all labeled cells of a lineage within the germarium. These are precisely the issues that were responsible for mistaken conclusions in the original FSC models and were addressed in the experiments and conclusions of Reilein et al., (2017). All of the results concerning sporadic contribution of FSCs to the FC of each new egg chamber, the low average FC contribution of single FSCs and the location of FSCs in more than one AP plane and distributed among all radial positions were thoroughly demonstrated for single-color lineage experiments in Reilein et al., (2017) and produced the same outcomes, as described in great detail, for multicolor lineage analysis. Some of those previously unpublished single-color images are shown here to document these findings directly.

It is entirely understandable that the very first model and description of FSCs in 1995, which represented a huge step forward (Margolis and Spradling, 1995), should be revised over time in light of better imaging and lineage tracing methods, together with a broader appreciation of possible stem cell organizations, most crucially involving the concept of population asymmetry (Jones, 2010; Snippert et al., 2010; Clevers and Watt, 2018). Rather than appreciating and further developing these revisions, for which overwhelming evidence has been presented in Reilein et al., (2017) and Reilein et al., (2018), and which were disseminated at Drosophila conferences as early as 2013, others have not cited or built on these crucial contributions but chosen instead to stick to old dogma (Cook et al., 2017; Kim-Yip and Nystul, 2018). We present a detailed account of the comprehensive shortcomings of the experiments and presentation of Fadiga and Nystul, and we explain why we stand firmly behind all of the conclusions made in Reilein et al., (2017).

Our intention is to make as clear as possible the overwhelming case that there are far more than two FSCs in each germarium (roughly 14-16), that the FSCs are present in more than one AP plane, that FSCs are maintained by population asymmetry, that single FSCs produce FCs only intermittently and that FSCs can produce ECs as well as FCs. It is crucially important for the wider community of stem cell researchers to be clear on our current understanding of these fundamental aspects of FSC organization and behavior, and the strength of the underlying evidence, despite efforts to portray these conclusions as controversial or incorrect, in order that discoveries made by studying FSCs can be widely appreciated and have an impact on our broader understanding of adult stem cells. Many of the issues we discuss, particularly with regard to lineage tracing and population asymmetry, are common to the investigation of many other types of adult stem cell, including the mouse intestine, where the organization of stem cells is in fact remarkably similar to that of FSCs (Snippert et al., 2010; Ritsma et al., 2014; Beumer and Clevers, 2016). We believe that detailed consideration of these issues with reference to FSCs should therefore be instructive and of interest to investigators studying any type of adult stem cell.

## Results

The Results section of Fadiga and Nystul is devoted to investigation of FSCs first by multicolor lineage analysis and then by single-color lineage analysis. Below, we consider these two subjects in sequence and then the conclusions that Fadiga and Nystul present in their Discussion section.

It is important to begin by defining FSCs clearly. Drosophila females produce eggs throughout adult life. The populations of germline and somatic cells that maintain a lifelong supply of the transient derivative cell types that contribute directly to the continued production of eggs are therefore termed stem cells. Eggs develop from a sequence of increasingly large egg chambers that first bud from the germarium at the anterior end of each ovariole (Fig. 1) (Duhart et al., 2017). An egg chamber contains germline cells surrounded by a monolayer of Follicle Cells, with specialized Polar Cells at either end and specialized Stalk Cells connecting neighboring egg chambers. The precursors of these somatic cells are sometimes referred to as pre-follicle cells in the germarium but here and in Reilein et al., (2017) we simplify the terminology by referring to all of these cells as Follicle Cells (FCs). Importantly, the defining feature of an FC is that it associates with a germline cyst, moves out of the germarium with that cyst and continues that association as the egg chamber matures. The somatic cells that are not themselves FCs but maintain production of FCs are called FSCs. Thus, FSCs are upstream of FCs. This defining difference is determined experimentally by timing (Fig. 1). A marked FC produces a lineage associated with a single cyst and must pass through the ovariole with that cyst (with known timing), producing a transient clone that passes through the ovariole within about five days (Margolis and Spradling, 1995). Any lineage mark (initiated in all experiments to be discussed at a defined time by heat-shock induction of a relatively short-lived recombinase) that is seen in FCs anywhere in an ovariole beyond this time window (more than five days after heat-shock) must have originated in an upstream cell, an FSC, and the lineage is therefore referred to as an FSC lineage (Fig. 1). This fundamental definition of the difference between an FC and FSC is consistent with the first definition of FSCs (Margolis and Spradling, 1995) and with a general principle that can be applied to all adult stem cells (Clevers and Watt, 2018; Post and Clevers, 2019); the definition must be applied rigorously and universally, no matter what is the lifetime of the initiating FSC or the frequency of FC contributions.

**Figure 1.**
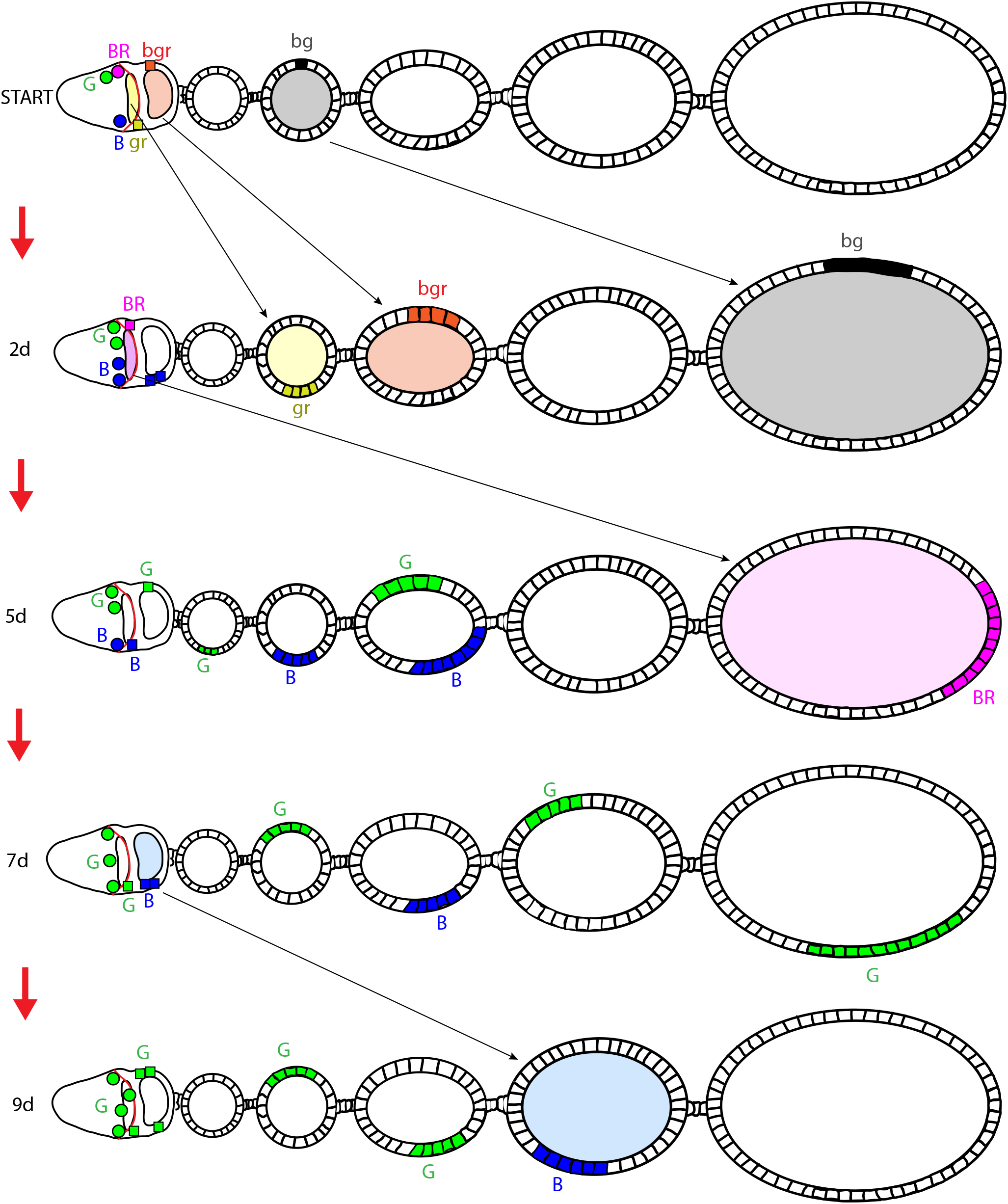
Temporal definition of a marked lineage initiated in a FSC. The cartoon illustrates the germarium (left), housing a lens-shaped stage 2b germline cyst (yellow), followed by a more spherical stage 3 germline cyst (orange) and five egg chambers of increasing size. Shortly after heat-shock (by about 0.5d and labeled “START”) a subset of proliferating FSCs and Follicle Cells (FCs) have undergone recombination and are depicted by six examples in different colors. By 2d, after about 3 cycles of egg chamber budding, all initially labeled cells associated with a germline cyst (yellow, orange and black squares) have amplified into a contiguous patch and have moved posteriorly (to the right) together with their associated germline cysts, which are shown with matching colors to illustrate continued association. The characteristic of stable association with a germline cyst defines a FC (including those in the germarium, which are sometimes called pre-follicle cells) and is the basis for depicting those cells with a square and a lower-case label. By 5d, even the earliest FC on a stage 2b cyst at 0.5d after heat-shock (yellow; gr), together with its entire lineage, has exited the ovariole (which extends beyond the five egg chambers shown), leaving no trace. Any labeled FCs present anywhere in the ovariole five or more days after heat-shock induced recombination must therefore derive from a recombination event in a cell upstream of an FC; all such cells are defined as FSCs (they are depicted here as circles with upper case labels). The behavior of individual FSCs is the subject of FSC lineage tracing experiments. It must therefore be considered unknown (no assumptions) at the start of an experiment (and it turns out to be stochastic). Three possible examples are illustrated. The magenta (BR; blue plus red) FSC survives for a couple of 12h cycles of egg chamber budding but it is then lost, becoming a FC on a stage 2b cyst (which is therefore a square colored a matching magenta color) by 2d. The lineage originating in the magenta (BR) FSC is still present as a single FC patch at 5d but it is lost from the ovariole by 9d. The blue (B) FSC survives longer, occasionally producing FCs (blue squares) but at some time between 5d and 7d the last blue FSC is lost. The last blue FC produced (at 6.5d) matures into a FC patch in the fourth egg chamber at 9d, with no other remnant of the blue lineage. The green (G) FSC initially produced no FCs but amplified and then green FSCs continued to produce FCs intermittently, while maintaining an FSC population that varies in number over time, with four green FSCs present at 9d. In an experiment, the only information available is the time at which cells were labeled by recombination and the appearance of labeled cells in the ovariole at 9d. Every labeled FC patch observed at 9d must have derived from an initially labeled cell that was not associated with a germline cyst (FSC circles at START). Every initially marked cell that was associated with a cyst (FC squares at START) had exited the ovariole, along with its entire lineage at least 4d earlier (by 5d). The 4d leeway is more than sufficient to be certain there is no possibility of slightly delayed recombination due to Flp recombinase perdurance allowing a FC lineage to remain at 9d (the FC would need to be labeled 4d after heat-shock). In the example illustrated, the green lineage would be scored as a FSC lineage with at least one surviving FSC. The blue lineage would be scored at 9d as an FSC lineage with no surviving FSC (and it can be inferred that there was a surviving blue FSC at 5d). Note also that the blue FSC lineage would only be detected if at least four egg chambers were scored. The purple lineage did originate in an FSC but it would be missed because the ovariole is scored at 9d (to leave no possibility of lineages originating from an FC) rather than after the minimal possible time of 5d. In other words, some FSC lineages are missed in order to be quite certain that all of those seen and scored did arise in an FSC. This experimental definition of a FSC does not rely on knowing or assuming where FSCs are located. The location of FSCs can best be determined by assessing the location of cells matching a FSC lineage color when it is the only such candidate cell in the germarium. Such analyses in Reilein et al., (2017) placed FSCs most frequently in “layer 1” immediately anterior to the anterior border of strong Fas3 staining (indicated by a red line along the posterior surface of stage 2b cysts), slightly less frequently in “layer 2” adjacent and anterior to layer 1 or less frequently in layer 3, adjacent to layer 2. The exact dynamics of somatic cell associations with cysts in the germarium are not known but are also not relevant for the definition of an FSC. If a cell initially associated with a cyst in the germarium but did not emerge with that cyst in a budded egg chamber, the lineage derived from that cell would, appropriately, be scored as originating in an FSC.

### Multicolor lineage Analysis

The number of FSCs that maintain FC production in an ovariole can, in principle, be determined by measuring the fraction of all FCs produced by a single FSC, by counting the number of FSC lineages in an ovariole (if each FSC lineage can be endowed with a different label) or by identifying the locations of all FSCs. All three approaches were used by Reilein et al., (2017). In each case, a large number of ovarioles must be sampled without any selection bias to calculate a reliable average and, crucially, to take account of any non-homogeneous behavior.

Reilein et al., (2017) devised a strategy to count the number of FSC lineages according to the loss of GFP, *lacZ* and RFP markers through FRT-mediated recombination events. In that system, recombination stochastically eliminates either GFP or *lacZ* (through recombination at *FRT40A* on the 2L chromosome arm) and, independently, may eliminate RFP (through recombination at *FRT42B* on the 2R chromosome arm), potentially generating five distinguishable recombinant genotypes in addition to the parental genotype. This system was chosen, and others rejected, because it demonstrated the key features of very low background recombination without heat-shock induction of the *hs-flp* gene encoding Flp recombinase and similar, tunable (by heat-shock severity) but potentially high rates of recombination at *FRT40A* and *FRT42B*. The low background without heat-shock is essential to deduce that experimental lineages resulted from recombination at a known time (shortly after heat-shock), and hence that all marked cells observed five or more days later derived from recombination in an FSC (a FSC lineage) (Fig. 1). The other key virtue is the ability to label six distinguishable lineages. In order to take advantage of this facility to visualize as many FSC lineages as possible in a single ovariole it is important that a high proportion of FSCs undergo recombination and that as much of the ovariole as practical is scanned for the presence of marked FCs. An ovariole displays a history of a limited period of FC production. Egg chambers generally bud every 12h, so scoring the FCs around six egg chambers and two germline cysts reveals about 4d of FC production at most (Fig. 1). Also, two or more FSCs in a given ovariole may sometimes undergo the same stochastic type of recombination (to produce the same combination of markers), so the number of distinguishable FSC lineages seen in an ovariole reports the presence of at least that number of FSC lineages but it is generally an underestimate. The underestimate will be severe if there are few colors relative to the number of FSCs or if some colors are adopted much less frequently than others (Fig. 2B).

**Figure 2.**
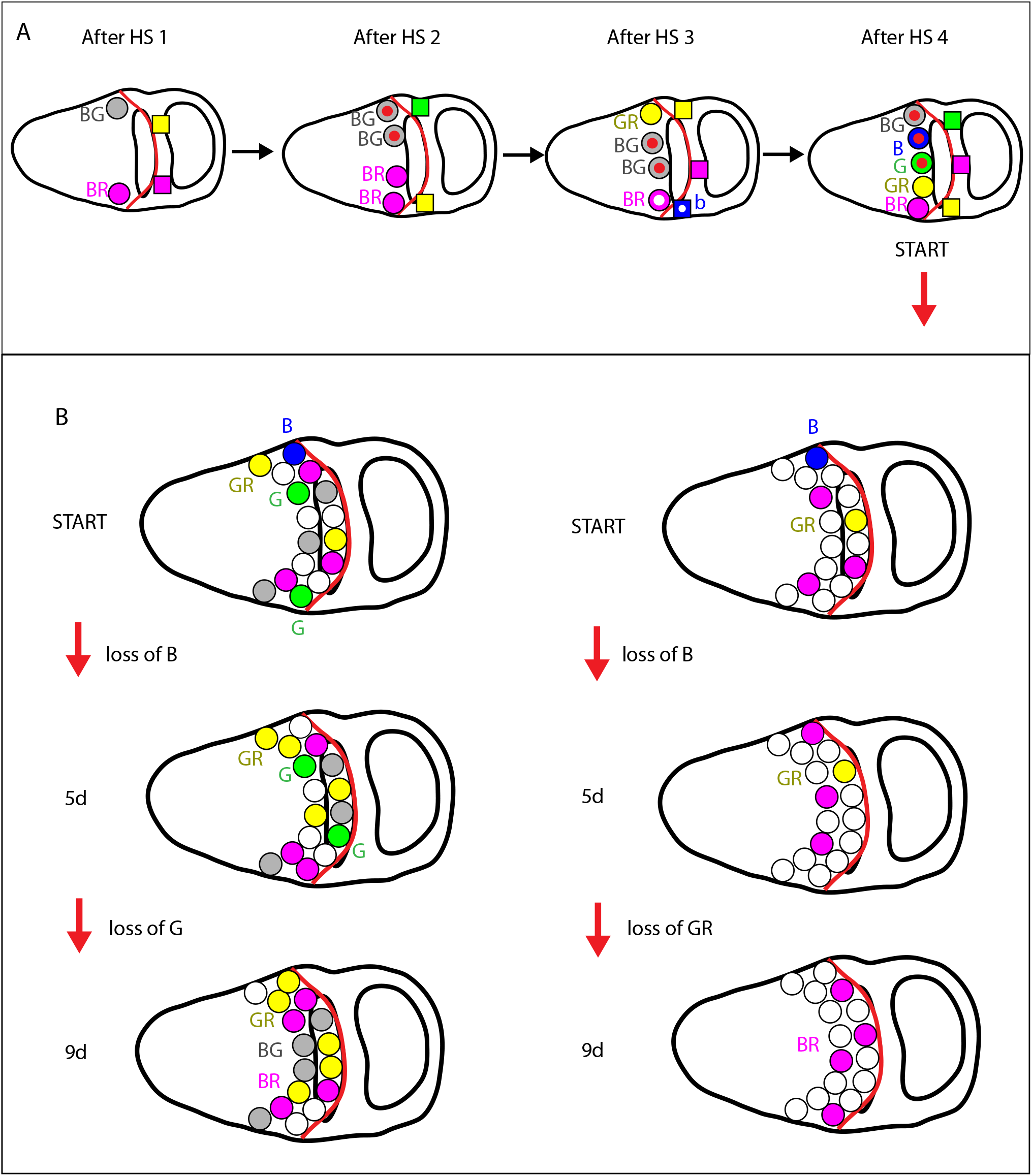
Multiple heat-shocks and the key role of high frequency marking of FSC lineages in multicolor lineage analysis for counting the number of FSC lineages. (A) The pictured germaria illustrate a possible sequence of events when using four spaced heatshocks (“HS”) to elicit four rounds of possible recombination events. As in Fig. 1, FSCs are shown as circles and the anterior border of strong Fas3 staining (red) is shown, close to the posterior surface of the stage 2b germline cyst. Fadiga and Nystul (2019) and a reviewer stated that the use of multiple heat-shocks is problematic because “subclones” are generated. A grey (BG) FSC generated by recombination shortly after the first heat-shock later duplicates and then one BG FSC undergoes recombination after the fourth and final heat-shock to produce B and G daughter FSCs (“subclones”, following the exact example cited by a reviewer). A red dot in the center of these cells indicates that they are related by lineage, as described. No further recombination takes place because there are no further heat-shocks and the timed experiment begins now (“START”), with ovarioles examined 9d later. Those ovarioles will show the presence of B, G and BG FSCs (among others) if those FSC lineages are not lost early. It will be deduced correctly that the B, G and BG lineages derived from an FSC of the corresponding color at the start of the experiment. That is equally true whether B and G are “subclones”, as shown here, or if they had no lineal relationship to each other or the BG FSC, as in the case of the illustrated GR (yellow) and BR (magenta) FSCs. Whether the cell has a red dot or not (indicating its past history) has no influence on how it will behave or its identity as an FSC. All FCs with recombinant genotypes that are produced at any stage during the period of multiple heat-shocks (indicated by squares, and showing only a few for simplicity), prior to the start of the experiment, will not be represented in ovarioles 9d later. That remains true whether single or multiple recombination events took place in the FCs themselves or if some took place in FSCs, even if the sequence of recombination events produces “subclones”, as illustrated by a shared white dot for the blue (b) FC generated from a magenta (BR) FSC. Again, the presence or absence of dots, indicating earlier lineage or “subclones”, is of no consequence; the only key issue is the identity of the cell as a FSC or FC (circle or square), which is revealed by the appearance of ovarioles 9d later, as explained in Fig. 1. (B) The pictured germaria illustrate typical plausible scenarios when recombination occurs at high frequency (left) or low frequency (right) among FSCs, corresponding to the conditions for Reilein et al., (2017) and Fadiga and Nystul (2019), respectively. As outlined in the Recombination Frequency section of Multicolor lineage analysis in the Results section, the data of Reilein et al., (2017) and Fadiga and Nystul (2019) conform roughly to frequencies of producing recombinant colors of 0.7 and 0.3 per cell, respectively. For p=0.7 (left) there will, on average, be 11 recombinant colors among the 16 pictured FSCs, with lower frequencies of RFP-negative colors (because *RFP/+* and *RFP/RFP* are both RFP-positive). An average frequency of recombinant colors of 0.3 (right) will, on average, produce 5 recombinant genotypes among the 16 FSCs, with a very low frequency of B and G genotypes, which can only be generated by two recombination events. It is therefore expected that all five recombinant FSC colors will usually be represented among an average of 11 recombinant FSCs for p= 0.7 (the parental color is the sixth and will also generally be present) and usually no more than three different recombinant colors (B, BR and GR are shown) will be represented among an average of five recombinant FSCs for p= 0.3. Moreover, each of the three colors will necessarily be represented by only a small number of cells (here B and GR have just one FSC). Over time, colors represented initially by one or a small number of FSCs are the most likely to disappear. Reilein et al., (2017) deduced that about 2/3 of initial FSC lineages are lost over 9d. The illustrated examples conform to these expectations. By 5d the initial single blue (B) FSC is lost in both scenarios (left and right). Between 5d and 9d, for p= 0.7 (left) the green (G) lineages, initially represented by 2 FSCs are lost, whereas yellow (GR) lineages, also initiated by two FSCs, are retained. For p= 0.3 (right) the yellow (GR) lineage, initiated by only one FSC (fewer than in the germarium on the left), is lost between 5d and 9d. In both cases the number of FSCs in other lineages changes stochastically (due to FSC division and FSC loss), with the same behavior pictured for the three magenta (BR) FSCs that were present initially in both cases. In 9d samples, assuming the whole ovariole is thoroughly examined, in the ovariole derived from high frequency FSC labeling (left) there will be four colors of FSC lineages with surviving FSCs (BG, BR, GR and the parental white lineage) and another FSC lineage color without a surviving FSC (G; scored also as a surviving FSC at 5d). For the ovariole derived from low frequency FSC labeling there are only two FSC lineage colors with surviving FSCs (BR and parental) and one additional FSC lineage color without a surviving FSC (GR), which would be missed by Fadiga and Nystul if the FSC was lost prior to 7d because ovarioles were not scored beyond 2-3 egg chambers (as illustrated in Fig. 1). These depicted scenarios and outcomes are commensurate with the observations of Reilein et al., (2017) that more than 50% of ovarioles had five or more colors of FSC lineages, and more than 40% had four or more colors of FSC lineages with surviving FSCs at 9d, whereas Fadiga and Nystul (2019) reported that most ovarioles have fewer than three FSC lineages. The cartoons illustrate that both sets of results are consistent with the presence of multiple FSCs (sixteen here) but that, crucially, the number of FSCs present can only be measured accurately if different FSCs are made visible by labeling most, or all of them with different colors and detecting the presence of differently colored lineage by examining the whole length of the ovariole.

Fadiga and Nystul report that in their hands the multicolor lineage analysis (which they term the LGR system) using Drosophila stocks that we built and provided is not reliable and gives different results from those reported in Reilein et al., (2017). Specifically, they report an inexplicably high frequency of background clones and other alleged artifacts. Neither high background nor artifacts were seen in Reilein et al., (2017). Even more important, Fadiga and Nystul use conditions of assaying only early portions of the ovariole and low recombination frequency that greatly diminish the potential to visualize multiple FSC clones.

#### (a) Background clones

Fadiga and Nystul report that in the absence of deliberate heat-shock 25% of ovarioles had recombinant clones that included an FSC and an additional 22% had clones affecting only FCs in 7d – old LGR flies. As they state, we reported a background clone rate of less than 2% in Reilein et al., 2017, over twenty times lower. Fadiga and Nystul use a control of *FRT19A* generating GFP-negative clones and report a background clone frequency of less than 2%. We have found *FRT19A* recombination efficiency to be consistently lower than for *FRT40A* (and other *FRTs* we have used extensively at 42D and 82B). In Fig. 4C, Fadiga and Nystul report an FSC clone frequency of less than 40% for the *FRT19A* experiment 7d after four 1h heat-shocks, while we consistently find a frequency in excess of 80% at 6d using *FRT40A* after a single 1h heat-shock at 37C (Melamed et al., unpubl.). Thus, as a minor point, the *FRT19A* control is a very insensitive “control”, allowing detection of only one type of recombination event (rather than three possible events for LGR) and uses an *FRT* with notably low frequency of recombination.

More than half of the background clones Fadiga and Nystul reported include FSCs (and many of the clones described as transient may also have originated in cells upstream of FCs (see section on single-color clones)). In adult ovarioles there are vastly more proliferating FCs than proliferating FSCs that could undergo a recombination event that would produce clones visible in the dissected 7d-old adults. Hence, it is very likely that most of the background clones reported without heat-shock were induced during development or even while sorting adults of the correct genotype, prior to putting experimental and “control” adults into the same incubator to control for inadvertent heat-shocks. There can be significant temperature elevations if sorting pads are illuminated at high intensity for long periods and for samples that are in the vicinity of, typically very hot, light sources for several minutes while sorting flies from other vials stored in the same container.

More important than the unknown cause of background clones in the experiment of Fadiga and Nystul, our no heat-shock controls, performed in parallel with our experimental heat-shock protocols showed an extremely low level of clone induction (Reilein et al., 2017). We took great pains to derive a multicolor system with this characteristic (testing several others along the way) and drew attention to this feature as providing us with the assurance that almost every one of the experimental clones analyzed was induced at the time of heat-shock (Reilein et al., 2017). The low background clone rate we found (and continue to find) is in keeping with the characteristics of the component *FRTs* at 40A and 42B, which consistently have similarly low background clone rates in isolation. It is hard to conceive of why combining the two should change that characteristic.

Fadiga and Nystul also present results for experimental clones, even though interpretation is clouded by their high background clone frequency. The most important issue concerns the relatively small number of FSC clones they report per ovariole (discussed in (d) below). They also report some additional problems with their analysis that we did not encounter (discussed in (b) below) and incorrectly describe some observations and assumptions in Reilein et al., (2017) concerning the frequency of different types of recombination events (discussed in (c) below).

#### (b) Clones lacking both 2L markers

None of the expected potential recombination events involving *FRT40A* should eliminate both the GFP and *lacZ* marker genes because these transgenes are heterozygous and both are situated on the left arm of chromosome 2 (2L) distal to *FRT40A* in LGR flies (see Fig. S1 and Fig. S7 in (Reilein et al., 2017). Fadiga and Nystul report detection of some FC patches lacking both GFP and *lacZ* signal but do not describe the frequency of such patches. They cite the presence of GFP and *lacZ* signal elsewhere in the ovariole as a control for marker detection. However, the GFP and *lacZ* signals are not equally strong throughout the ovariole and the quality of antibody staining and imaging is crucially important. For the examples shown, which are presumably the clearest available, the intensity of *lacZ* staining in neighboring cells appears to be both low and variable in Fig. 2C and 2E, and the same is true of GFP staining in Fig. 2D. Thus, the evidence for the reported unexpected staining patterns originating from genetic or epigenetic changes does not appear convincing; imperfect antibody staining is a more plausible explanation.

More important, we had ample opportunity to see any such exceptions in our analyses because we spent hundreds of hours scoring every cell in every z-section of such samples. We noticed that the *lacZ* marker was generally expressed at lower levels in later egg chambers and that the exact staining protocol needed to be carefully optimized in order to score all cells, but we did not find FC patches lacking both GFP and *lacZ* or any other unexpected phenotype. Thus, the problem Fadiga and Nystul cite was not evident in our studies (Reilein et al., 2017), is of unreported prevalence in their study and seems likely to result from deficiencies in experimental execution rather than any (unidentified) intrinsic defect in the LGR clonal system.

#### (c) Relative frequency of clone types

Fadiga and Nystul wrote that “An assumption that was integral to the calculations of FSC number in the Reilein et al. study was that the clone induction protocol induced recombination at both FRT sites in every mitotic follicle cell” (they presumably really mean FSC, as only that is relevant). That statement is incorrect on three counts.

First, the observed proportion of different clone types was used only in a very minor way towards calculating FSC numbers. When we counted the number of different FSC clone types with a surviving FSC at 9d after heat-shock we saw a distribution that is shown in Fig. 2i of Reilein et al., (2017) and Fig. 1E of Fadiga and Nystul (2019). The number of different FSC clones in a single ovariole generally underestimates the actual number of lineages because two or more FSCs can undergo the same type of recombination yielding two or more lineages of the same color. Some type of statistical inference is therefore needed to use the raw data of the observed number of different types of FSC clone in each ovariole to derive an estimation of the average number of surviving FSC lineages among all scored ovarioles. We used a method, outlined below, to estimate that there was an average of about five surviving FSC lineages at 9d after heat-shock. No matter what statistical inferences are used it is clear from the raw distribution of distinguishable FSC clones (Fig. 2i of Reilein et al., (2017) and Fig. 1E of Fadiga and Nystul (2019)) that the average number of surviving FSC lineages could not possibly be much lower than five because almost half of the ovarioles had at least four differently colored lineages. Thus, the impact of the statistical method chosen is extremely minor, not “integral” or at all significant in the context of whether there are two FSCs or fourteen to sixteen in total.

Second, we reported that the experimental distribution of clone types we observed approximately matched the outcome that would result from a simple theoretical scenario that we described (where the probability of recombination was 2/3 at *FRT40A* and was also 2/3 and independent at *FRT42B* as a final outcome of the multiple heat-shocks, considering this as a single process, leading to a 1:1:1:2:2:2 frequency of the six possible color combinations). We summarized the experimental data in the Supplementary Note by saying that “the three types of RFP-negative FSC clones (B, G and BG) were present at roughly 1/9 [each] of the total, while the three types of RFP-positive FSC clones (BR, GR, BGR) were present at a frequency of roughly 2/9” (B stands for *lacZ* visualized in Blue, G for GFP and R for RFP). The actual numbers for FSC clones with a surviving FSC were 13, 19, 17, 38, 44 and 40 for B, G, BG, BR, GR and BGR, respectively. We did not start by assuming a certain hypothetical scenario, as Fadiga and Nystul imply; we first observed a specific distribution of clone types and noticed that, by good fortune, the outcome matched a very simple model that allowed a simple statistical approach, which is fully documented in the Supplementary Note.

Third, Fadiga and Nystul imply that (i) Reilein et al., (2017) assume there is recombination at each FRT in every FSC and (ii) that this is not possible, *inter alia*, because cells must be in a specific phase of the cell cycle.

i. We neither modeled nor reported results consistent with recombination at each *FRT* in every cell. The model has a net recombination frequency of 2/3 at each site (above) and experimentally about 2/9 of clones retained all three markers (no recombination at either FRT site). Experimentally, we observed a nearly even distribution of 2L genotypes (*GFP/GFP, GFP/lacZ, lacZ/lacZ*) and that RFP on 2R was absent in about one third of clones, consistent with a roughly equal representation of the possible 2R genotypes (+/+, *RFP/+*, *RFP/RFP*) and a similar distribution of 2L genotypes for RFP-positive and RFP-negative clones. If there had been recombination at each *FRT* in every cell there would instead be no clones with both GFP and *lacZ*, and half of the clones, not a third, would lack RFP. Thus, the explanation that Fadiga and Nystul offer in the main text for the results that we observed is quite incorrect and the “predicted” distribution of clone types shown in Fig. 3A of Fadiga and Nystul is not what would result from the model they describe in the Figure legend.
ii. The experiment involved four heat-shocks. It is therefore quite reasonable that each FSC was at a suitable stage of the cell cycle during one or more heat-shocks. Also, the perdurance of the *hs-flp* Recombinase product is not known well enough in these or any other cells to know what range of cell cycle positions at the time of heat-shock permits recombination (Beumer et al., 1998). From these two considerations it is not possible to predict what is the maximum *FRT*-mediated recombination frequency that is theoretically possible. It is also, of course, quite reasonable that we achieved at least one recombination event in at least 7/9 of all FSCs, consistent with the experimental observation that about 2/9 retained all three parental markers.

**Figure 3.**
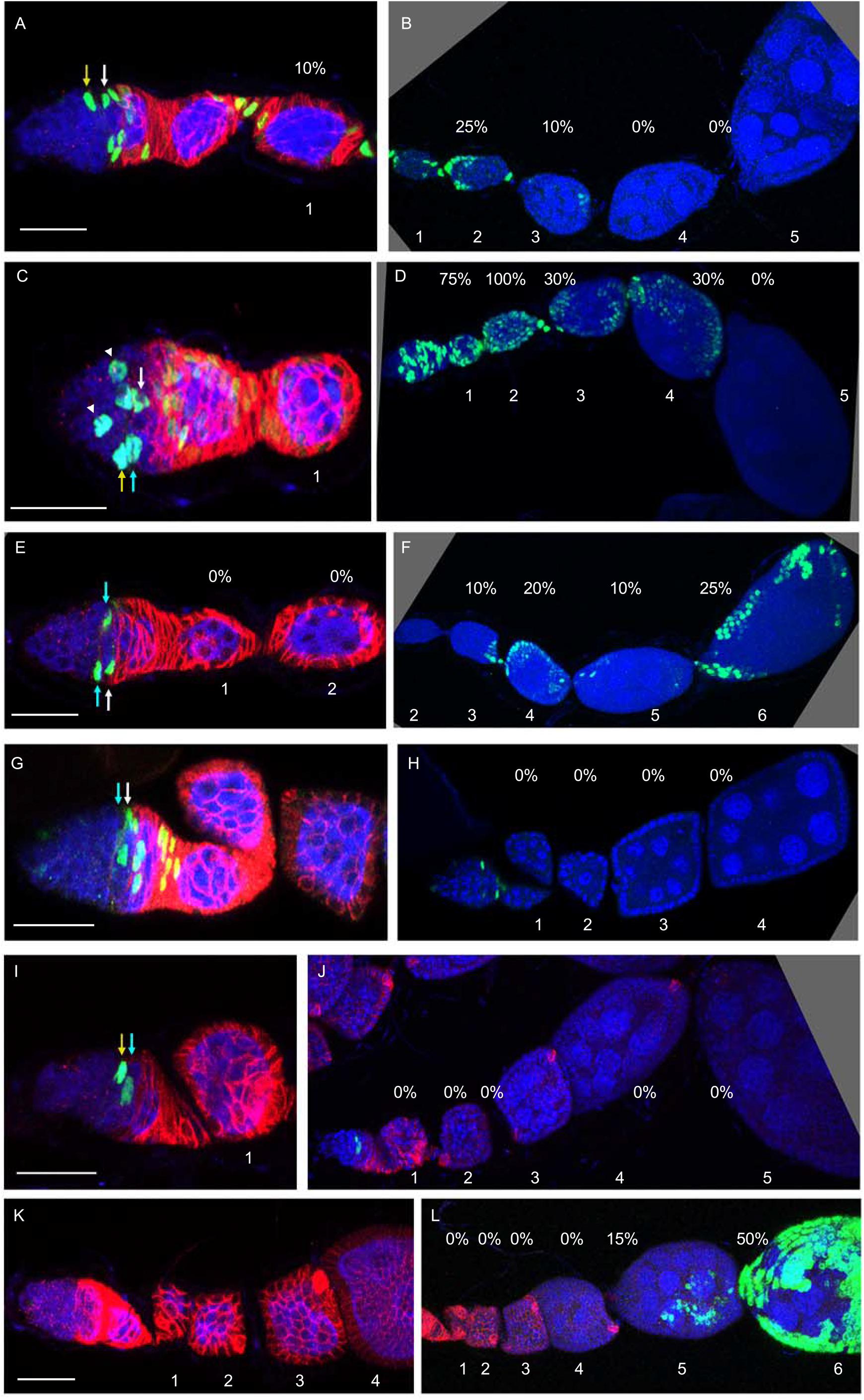
MARCM FSC lineages illustrate the heterogeneity of FSC behaviors, frequent FSC loss and FSC amplification. (A-L) A selection of MARCM clones from a single experiment examining ovarioles 9d after clone induction. Panels on the left include the germarium and early egg chambers, while panels on the right show the remainder of the same ovarioles; egg chambers are numbered to make clear overlapping regions. GFP-labeled cells (green) in each lineage are shown together with DAPI staining of all nuclei (blue) and Fas3 antibody staining (red) in select panels. Each image includes only a limited set of z-sections, so not all GFP-positive cells are visible. FSCs are indicated by arrow colors as being in layer 1 (white), layer 2 (blue) or layer 3 (yellow). The estimated proportion of the FC epithelium of each egg chamber that has GFP label is shown by the indicated percentage values (calculated by looking through all z-stacks). The final average value of 15% labeled FCs was calculated from all lineages (over sixty) with at least one marked FSC and FC patch (Reilein et al., 2017). (A-D) Ovarioles with marked FSCs and FCs in both the germarium and one or more egg chambers (36/148 ovarioles; 120/186 egg chambers with marked FCs; average of 4.4 FSCs). (E, F) An ovariole with marked FSCs and FCs in one or more egg chamber but not in the germarium (16/148 ovarioles; 50/84 egg chambers with marked FCs; average of 2.1 FSCs). (G, H) An ovariole with marked FSCs and FCs in the germarium but not in any egg chamber (10/148 ovarioles; average of 4.4 FSCs). (I, J) An ovariole with marked FSCs but no FCs (21/148 ovarioles; average of 1.7 FSCs). (K, L) An ovariole with marked FCs but no FSC (14/148 ovarioles). Several of the lineage categories above also included ECs. Remaining ovarioles (not shown) included ECs but no FSC or FC (20/148) or no marked cells (31/148). Only 24 lineages included marked FCs in the germarium and both of the first two egg chambers (the criterion used by Fadiga and Nystul for selecting lineages to score FC contributions) out of a total of 117 ovarioles with marked cells, of which 76 are definitive FSC lineages judged by the presence of marked FCs.

To summarize, the protocol we used, partly by good fortune, produced a highly desirable outcome where the representation of all possible genotypes was roughly equal (and RFP-positive phenotypes are twice as common as RFP-negative phenotypes because they are represented by twice as many genotypes: *RFP/+* and *RFP/RFP*). The most important consequence of this distribution was that it conferred high sensitivity for detecting distinguishable FSC lineages. A secondary, rather trivial consequence was that it also allowed us to use a very simple statistical approach to make a best estimate of the average number of surviving FSC lineages from the raw data, essentially accounting for separate lineages that could not be distinguished because they are the same color.

#### (d) Numbers and types of FSC clones

Fadiga and Nystul report, in Fig. 1D and 1E, a much lower number of different FSC clone colors per ovariole than we found using the multicolor, LGR, lineage analysis. This is most pronounced and relevant at relatively early times after heat-shock, including 9d. This difference could theoretically have several origins, which are considered separately below.

##### (i) Scoring

In Reilein et al., (2017) we carefully described and showed exactly what we were scoring. We captured, stored, and scored all z-sections in each color channel (for GFP, *lacZ*, RFP, Fas3) through most of the length of each ovariole, together with images of each germarium at higher magnification so we could score the precise location of all potential FSCs, as illustrated by the samples shown in Fig. 1, Fig. 2 and Figs. S1-S3 of Reilein et al., (2017). We noted that even a single FC patch found at 9d after heat-shock must have resulted from recombination in an FSC and therefore reveals an FSC lineage. We also looked for a candidate FSC of the same color in the germarium and if there was at least one such cell we scored that as an “FSC lineage with a surviving FSC”. We reported the number of distinguishable (by marker color combination) FSC lineages with a surviving FSC in each of 50 ovarioles (graphed in Fig. 2i and tabulated in the Supplementary Note of Reilein et al., (2017)).

Fadiga and Nystul did not describe exactly what they are scoring in the main text (Fadiga and Nystul, 2019). The legend to Fig. 1 says “(A-G) Analysis of LGR clones within the follicle epithelium, ranging from the Region 2a/2b border of the germarium to the first 2-3 follicles downstream from the germarium….” and Fig. 1 D, E and G report the numbers of differently colored clones per ovariole. The documented e-Life reviewer exchange confirms the description in the Figure legend that ovarioles were only scored in their anterior regions. By failing to include egg chambers beyond the first two or three, the authors missed a significant segment of crucial data and therefore undoubtedly underestimated the number of FSC lineages present (see Fig. 1 for an illustrated example). In Reilein et al., (2017) we were able to score five egg chambers for 8 ovarioles, four egg chambers for 28 ovarioles and three egg chambers for 12 ovarioles among the 50 ovarioles reported 9d after heat-shock. The total number of FSC clones with a surviving FSC was 171 (an average of 3.4 per ovariole). Re-examination of the spreadsheet of all clone locations allows us to evaluate how many FSC lineages we would have missed if we had scored fewer egg chambers. Had ovarioles been scored only up to egg chamber 3 (even that is apparently more than scored in some ovarioles of Fadiga and Nystul), we would have missed ten FSC clones with surviving FSCs and we would have missed sixteen such clones had egg chamber 3 also not been included. If we consider all FSC clones (those with labeled FCs, whether there is a surviving FSC or not) we would have missed nineteen clones by scoring only three egg chambers and thirty-two had egg chamber 3 also not been included. For stem cells maintained by population asymmetry the highest rate of loss of lineages is at the earliest times (and the derivatives of FSCs lost early would appear only in the most mature egg chambers of dissected ovarioles); later, most surviving stem cells have duplicated, reducing the rate of lineage loss. It is therefore extremely important to assay as much of each ovariole as possible to maximize sensitivity of detection of FSC lineages. The purpose of the multicolor lineage experiment is to visualize as high a proportion of the FSC lineages present as possible, requiring maximal sensitivity in all facets of its application.

##### (ii) Recombination efficiency

Even taking into account the above consideration of missing several FSC clones, the number of distinguishable clones Fadiga and Nystul report is still considerably lower than observed by Reilein et al., (2017). The key consideration is whether this results from a genuinely smaller number of FSCs that could potentially acquire different lineage colors or if there was insufficient opportunity for different FSC lineages to be revealed because recombination frequencies were too low. The frequency of ovarioles with no recombinant genotypes clearly shows that the latter explanation is correct. The evidence that Fadiga and Nystul induced recombinant clones at a much lower frequency than in Reilein et al., (2017) is clearest for the single heat-shock results. Fadiga and Nystul report in Fig. 1G a frequency of ovarioles with only a single lineage color, which is presumably in all, or almost all, cases the parental color prior to any recombination of over 50% at 9d and over 60% at 14d after a single 1h 37C heatshock. The corresponding frequency cited in the Fig. 2 legend of Reilein et al (2017) was 11% for the equivalent experiment in Reilein et al (2017) 12d after a single 1h 37C heat-shock. For the same heatshock conditions we have conducted nine experiments in the last five years using single-color MARCM at *FRT40A* and found an average frequency of ovarioles with no FSC clones of 18% at 6d and 26% at 12d. For the MARCM clones using *FRT40A* reported in Reilein et al (2017) 9d after a shorter, 15 min heat-shock at 37C the frequency of ovarioles with no FSC clones was 20% (31/148). Thus, the frequency of *FRT*-mediated clone induction we found using the LGR system is consistent with large numbers of our experiments using one of the LGR sites (*FRT40A*) and the same *hs-flp* transgene. This rate is much higher than Fadiga and Nystul report for the LGR flies, suggesting a deficiency in the flies or methodology that they used. That deficiency is important because it severely limits the ability to visualize a significant fraction of all of the FSCs present as distinguishable FSC lineages (Fig. 2B). The maximal number of differently colored FSC lineages that can be generated cannot be determined by an experiment that fails to provide the majority of FSCs with an opportunity to generate a recombinant FSC lineage.

The impact of recombination frequency is illustrated by considering different specific values of recombination frequency (illustrated in Fig. 2B). Specifically, we consider here a recombination frequency, p= 0.3 (which likely roughly matches that seen for multiple heat-shocks by Fadiga and Nystul) and p=0.7 (which roughly matches the frequency seen in Reilein et al (2017). Imagine that there are on average five FSC lineages with surviving FSCs at 9d, as inferred by Reilein et al (2017). The probability of finding no recombinants (assuming independent events) is (1-p)^5^, which is 0.17 for p= 0.3 (similar to the value reported in Fadiga and Nystul (2019) for ovarioles with only one color at 9 dphs) and 0.002 for p= 0.7 (Reilein et al., (2017) found one example out of 50). Thus, the values of p=0.3 and p=0.7 are roughly consistent with available data from both experiments. To detect five differently colored lineages in an ovariole requires that at least four of the five FSCs initiating the surviving lineages underwent recombination and that they include at least four different recombinant colors. The probability of four or five recombinants of any color among five cells is 5p^4^(1-p) + p^5^ = 0.029 for p= 0.3 and 0.53 for p=0.7. This very large difference is amplified further when multiplying by the probability that at least four recombinant colors are different, allowing five colors in total, because the representation of different colors is more equal and more diverse for higher recombination rates (compare “predicted” and “9 dphs” columns in Fig. 3A of Fadiga and Nystul (2019)). Thus, even for p= 0.7 it is rare to detect all five surviving FSC lineages as having different colors (observed in 4% of ovarioles in Reilein et al., (2017)) and it would be much more than twenty-fold less likely under the approximate conditions of Fadiga and Nystul where p= 0.3. Even for recovery of at least three recombinant genotypes among five FSCs (allowing the possibility of detecting four or more colors in total) the probability (p^5^ + 5p^4^(1-p) + 10p^3^(1-p)^2^) for p=0.3 (0.16) is five-fold lower than for p=0.7 (0.84). Reilein et al., (2017) found four different colors or more at a frequency of 0.42, roughly consistent with a p value of 0.7 if roughly half of ovarioles with at least three recombinant genotypes included at least four different colors in total. The latter proportion would be several-fold lower for an experiment with the much lower diversity and less equal representation of recombinant colors seen by Fadiga and Nystul, so the proportion of ovarioles with four or more distinguishable colors would be expected to be much less than half of 0.16 if p= 0.3, roughly consistent with their observation of just one such ovariole among 43. This numerical example, roughly in keeping with the results from the two studies, illustrates the critical importance of a high recombination frequency and the availability of many recombinant colors in order to detect the presence of multiple FSC lineages. The large quantitative impact of these factors may not be intuitively obvious without considering numerical details explicitly (Fig. 2B).

The origin of the insufficient frequency of clone induction reported in Fadiga and Nystul is uncertain but there are some plausible possibilities. They report conducting heat-shocks in empty vials, whereas we use vials with food and therefore less air space. Our conditions certainly transmit temperature changes well because we routinely find that heat-shocks as short as 10 mins in a 32C water-bath are effective at eliciting *FRT*-mediated recombination. All of the FRT sites under discussion consist of a pair of *FRT* units and it is commonly known that efficacy can be reduced if one copy is eliminated by recombination between the paired *FRT* sites in parental stocks (evident from loss of the *w+* marker for *FRT42B*, but not for the tandem *FRT* sites of *FRT40A*) (Golic and Lindquist, 1989; Xu and Rubin, 1993). Consequently, stocks with *FRTs* are generally maintained in the absence of any *flp* recombinase gene. In the LGR system one parent does include the *hs-flp* transgene. We carry several isolates of the parental stocks. Subsequent to the studies reported in Reilein et al., (2017) we have used multicolor clonal analysis in other studies. On two occasions we did not recover as high a proportion of the rare clone types (*lacZ* only and GFP only, requiring recombination at *FRT40A* and at *FRT42B*) as we wished. We solved that problem each time by switching to a different parental stock isolate, which we now maintain in more than one stock. The several multicolor lineage experiments reported in Reilein et al., (2017) were performed between 2012 and 2016, we sent stocks to the Nystul lab in early 2017 and we most recently switched our use of parental stocks in 2019.

It is extremely important to generate FSC recombination at high frequency in order to have a sensitive test of the number of differently labeled FSC lineages (Fig. 2B). Fadiga and Nystul reported much lower rates of *FRT*-mediated recombination than achieved in Reilein et al., (2017). That is potentially attributable to deteriorating *FRTs* or an inefficient heat-shock protocol, and partly to missing some FSC lineages because only anterior regions of the ovarioles were scored. The net result is an inability to estimate the true number of FSC lineages that can be generated in a single ovariole. The failure to generate and detect marked FSC lineages of any recombinant genotype at high frequency is not evidence for the absence of many FSCs and their potentially differentially marked lineages, which were generated and counted at high frequency in Reilein et al., (2017).

##### (iv) Multiple heat-shocks

Fadiga and Nystul contest the idea that any patch of FCs with a recombinant genotype that is observed 9d or more after clone induction should be scored as an FSC lineage, even if there are no other cells of that genotype. As explained carefully in the Introduction and in Reilein et al., (2017), such patches must have derived from recombination in an FSC based on timing because all derivatives of FCs downstream of an FSC would have exited the ovariole (Fig. 1). This timing criterion is essentially THE definition of an FSC; 9d is a full 4d more than is strictly necessary to ensure a FSC origin, leaving no room for any reservations concerning Flp recombinase perdurance after heat-shock or plausible variations of normal developmental timing. Knowledge of timing depends critically on all clones being induced by the timed heat-shock, which was demonstrated unequivocally in Reilein et al., (2017) (see “Background clones” above).

Fadiga and Nystul claim a further problem with this designation of a lineage arising from recombination in an FSC by saying “it is generally inappropriate to use multiple heat shocks… because the later heat shocks may produce subclones within clones that were induced by the earlier heat shock treatments.” This statement apparently derives from reviewer comments communicated, partly in upper-case, by the Reviewing Editor. It is categorically incorrect. First and most important, the number of heat shocks is of no conceptual consequence (Fig. 2A). Their purpose is to produce a variety of genotypes in the FSC population and, in experiments with multiple heat-shocks there will indeed often be two separate productive recombination events (there can only be two, as loss of a marker on 2L or 2R is irreversible). The cell lineage experiment starts when all of the heat shocks and recombination events are finished; it tests what happens to the variously marked FSCs over the next several days. It is of no consequence whether two or more of the FSCs at the start of the experiment were labeled at the first, second, third or fourth heat-shock or are lineally related to each other during the period of administering heat-shocks; it certainly does not alter the fact that an FC patch identified 9d later must have arisen from a cell upstream of all FCs, which defines an FSC (Fig. 2A). Thus, the presence of a parental recombinant genotype and of a different recombinant derivative (“subclone”) in the FSC population is not in any way problematic. It is an important practical means to generate a diversity of FSC lineages. If the parental genotype or the “subclone” genotype or both are produced in FCs during the period of heat-shocks, whether they were first produced in FSCs or not, is also of no relevance because all of these FCs and their derivatives will have exited the ovariole at least 4 days before the ovarioles are examined.

Second, Reilein et al., (2017) also examined lineages resulting from a single heat-shock and found up to five differently colored lineages per ovariole at 12d after heat-shock, with an average of 2.6 (116 in 45 ovarioles). That value is lower than for multiple heat shocks examined at 9d (171 clones in 50 ovarioles; average 3.4) in Reilein et al., (2017) but much higher than the number of distinct clones reported for the experiment of Fadiga and Nystul analyzed 9d or 14d after a single heat-shock (shown in their Fig. 1G as 55 lineages among 36 ovarioles and 29 lineages among 22 ovarioles, respectively). These results demonstrate experimentally that the observation of multiple distinguishable FSC lineages in a single ovariole is not dependent on multiple heat-shocks, as is already quite clear from the theoretical considerations discussed above. These results also emphasize the importance of inducing a high frequency of recombination to optimize sensitivity for detecting multiple FSC clones. That is precisely why multiple heat-shocks were used, with no associated logical or practical detriment, in Reilein et al., (2017).

To summarize, Reilein et al., (2017) used the multicolor lineage system with high sensitivity to detect different FSC clones, they clearly described and justified the scoring of fully presented, high quality samples, as clones deriving from recombination in an FSC and whether a corresponding FSC still survived, and showed in the same experiments that there were virtually no clones in samples without heat-shock nor any significant aberrant or unexpected staining patterns. The use of multiple, effective heat-shocks was important to maximize FSC clone color diversity, as was scoring as large a fraction of each ovariole as possible, facilitating the estimation of the true number of FSC lineages present; broadly similar results were found also when using a single heat-shock. Fadiga and Nystul present a variety of criticisms of methodology and interpretation, all of which are misplaced, as described above. They also provide experimental data presented as faulting the LGR multicolor genetic system, reporting a high frequency of background clones, relatively low frequency of experimental clones and some unexplained staining patterns in their hands. The source of these problems in the Fadiga and Nystul experiments is not demonstrated and remains unclear. It is clearly not an intrinsic feature of the LGR multicolor lineage system that we developed and used with great success (very low background clone rate, very high experimental clone rate, no significant artifacts). Thus, we strongly disagree with every facet of their compound summary sentence, “These findings challenge the underlying assumptions of the analysis in the Reilein et al. study, and suggest that, even if the LGR system were reliable, the interpretation of the data leading to their conclusions about the number of FSCs per germarium are also flawed.” The LGR system was, and has continued to be reliable in our hands, providing the best approach possible to date for counting FSC lineages according to the defining functional criterion of being upstream of FCs.

### Single-color clonal analysis

The majority of FSC lineage analyses in the literature have used only a single marker. That approach does not offer the possibility of directly counting the number of FSC clones in an ovariole but it does allow characterization of the patterns of FSC contribution to FCs and other cells (here ECs are relevant), measurement of the proportion of all FCs in an ovariole that are labeled as a means to estimate FSC number, and observation of FSC locations. However, there are important potential shortcomings regarding these applications. One key consideration is how many FSCs are labeled in an ovariole. More than one FSC may be labeled either because more than one FSC initially underwent recombination or because an initially single labeled FSC produced additional labeled FSCs in the interval prior to assay. The first issue can be controlled to some degree by using a weak heat-shock to induce clones at low frequency. That remedy can be made more effective by using two FRTs in the multicolor LGR system and scoring only lineages with two specific recombination events. The second issue has often been overlooked (the term “FSC clone” superficially suggests the contributions from a single FSC) but it has a very large impact for FSCs.

The failure to appreciate that individual FSC lineages might be lost or amplify at high rates (characteristic of population asymmetry, as we found) underlies the mistaken assumption in the earliest characterization of FSCs, and now repeated by Fadiga and Nystul, that an FSC clone with an identifiable candidate FSC typically, or always, shows the output of a single FSC. It does not. Many ovarioles that initially included a labeled FSC have lost all labeled FSCs (or even all traces of the FSC lineage), while in many other ovarioles an initially single labeled FSC has amplified to produce several labeled FSCs. The essential experimental approach is to make no assumption about whether FSCs are frequently lost or amplify or not, but to try to score every ovariole that initially included a labeled FSC, regardless of its appearance at the time of assay several days later.

The second key consideration regarding the location of all FSCs in the germarium, and the separate concept of identifying the cells that maintain Escort Cell (EC) production during adult life, is that the method for marking lineages must allow clear scoring of all cells in the germarium. For this objective, positive marking of nuclei with GFP (MARCM) (Lee and Luo, 2001) is far superior to any negative-marking approach, whether single- or multi-color. We therefore used MARCM to verify the results of multicolor lineage analyses in Reilein et al., (2017) with regard to FSC clone patterns, FC contributions of an average FSC lineage, location of FSCs, and production of ECs from FSCs.

Fadiga and Nystul principally used GFP-loss clonal analysis with *FRT19A* to report the patterns of FSC clones and in an effort to calculate the average FC contribution of a single FSC. This was supplemented by using a small number of GFP-positive MARCM clones to measure FC contributions of FSCs. In both cases the reported results are strongly biased by deliberately measuring only the subset of clones likely to contain multiple FSCs and ignoring a large fraction of ovarioles in which FSCs were initially labeled. Fadiga and Nystul then used clones marked by loss of GFP to report on FSC location. That method of clonal marking has severe limitations for assaying all germarial cells. There is indeed no evidence that all marked cell locations in the germarium were scored and recorded from that analysis (or MARCM or multicolor lineage analysis), in contrast to the multicolor clonal analysis and MARCM analysis of Reilein et al., (2017). Consequently, Fadiga and Nystul are unable to assess how many FSCs are present in each lineage and where all of those FSCs are located. Fadiga and Nystul do not address EC production by FSCs, even though that is relevant to their analysis of FSC location, preferring to quote a conclusion suggesting separate EC and FC lineages (Kirilly et al., 2011) that was overturned in Reilein et al.,(2017). Consequently, Fadiga and Nystul ignored any labeled FSCs that they did detect in more anterior locations than their preferred designation.

#### (i) Pattern of FSC clonal derivatives

Fadiga and Nystul show some examples and summary numbers for the types of single-color clones they observed in Fig. 3B-G but they do not state what were the associated conditions of heatshock (which would affect the number of FSC clones per ovariole), or the time after heat-shock, which is crucial to determining the nature of clones.

If we assume that the time after heat-shock is at least 7d (since that is true for all of their other reported investigations of single-color clones), then all clones observed in FCs derive from recombination in an FSC (because FCs are by definition associated with germline cysts and all stage 2b cysts and later have exited the ovariole before 7d). Thus, clones that the authors term “transient”, as shown in Fig. 3D and graphed in Fig. 3B of their study, are in fact FSC clones (clones originating in an FSC). The logic is the same for clones labeled as “FSC loss” and “Small”; all originated in FSCs. Fadiga and Nystul presumably designate the clone in Fig. 3D as “transient” and the clone pictured in Fig. 3E as “FSC loss” because they are making the assumption that an FSC clone must be contiguous. It is obviously not reasonable to test whether FSC clones are always contiguous by first assuming that this is the case in order to score or classify the data. All of the types of clone shown, and given five different names, in Fig. 3B-G of that study, originated in an FSC.

Thus, the data summarized in Fig. 3B of Fadiga and Nystul (2019) show that at least 28.9% of FSC clones (“Transient” plus “Small”) included just a single FC patch. A further 9.6% (“FSC loss” and “Discontinuous”) had more than one FC patch but also had egg chambers or germarial regions with no labeled FCs. The remaining 61.5% of clones are labeled as “Normal” and are described as including labeled FCs in the germarium and in one or more egg chambers, implying that some unspecified proportion of egg chambers contain no labeled FCs. These data were aimed at investigating the quoted result from Reilein et al., (2017) that “each FSC does not normally contribute to every follicle”. The data in Fig. 3B, when described properly, support that statement (38.5% plus some unknown fraction of 61.5% show that the initially labeled FSC did not contribute to every follicle), yet the authors introduce these data by saying that “the clone patterns in 94% of ovarioles with clones did not fit this expectation”. That statement is a flagrant mis-representation of the data.

If we were to score the data from Reilein et al., (2017) for clones examined 9d after induction according to the quoted criterion for “Normal” in Fig. 3B (labeled FCs in the germarium and at least one egg chamber) the proportion of such clones among all FSC clones (those with any labeled FCs) is 61% (130/211) for the multicolor lineage analysis and 66% (51/77) for MARCM clones. These values are very similar to the 61.5% reported by Fadiga and Nystul, contrary to their assertion that the distribution of FSC clone types is very different in the two studies. Only the description of the data differs for this particular characteristic. Of the clones from Reilein et al., (2017) that would fit the description of “Normal” used by Fadiga and Nystul the proportion that contribute to every scored egg chamber is 32% (41/130) for the multicolor lineage analysis (which scored 5.3 egg chambers per ovariole on average) and 20% (10/51) for the MARCM analysis (which scored 3.8 egg chambers per ovariole on average). Thus, contribution of FCs to the germarium and each egg chamber is unusual among FSC clones, especially when many egg chambers are scored (41/211 or 19% when scoring an average of 3.8 egg chambers in the multicolor analysis, and 10/77 or 13% when scoring an average of 5.3 egg chambers in the MARCM analysis).

Thus, the GFP-negative clonal data presented by Fadiga and Nystul are described with misleading labels but actually support the observation of Reilein et al., (2017) that the majority of FSC clones do not include FCs in every egg chamber. Importantly, Reilein et al., (2017) used appropriate logic and terminology, and found consistent results using both LGR multicolor labeling and MARCM.

#### (ii) Proportion of FCs contributed by a single FSC

The exact method for trying to measure the proportion of all FCs in an ovariole contributed by a single FSC, together with the associated limitations, are extremely important to consider carefully in order for the result to be unbiased and allow an estimation of the contribution of a single FSC. Fadiga and Nystul do not describe their method in detail, but write in the Methods: “To determine FSC clone sizes, ovarioles with clones that extended from the Region 2a/2b border to at least a Stage 2 follicle were selected for analysis.” It is inappropriate to introduce any bias from selective sampling and the criterion used by Fadiga and Nystul introduces a very strong bias. The appropriate, correct approach is (as far as possible) to include all ovarioles in which an FSC was originally marked, no matter what the appearance of the clonal derivatives. The problem of bias in sample selection would be further compounded if the authors counted only, or even predominantly, the part of each ovariole where FC presence is enriched by sample choice (germarium and early egg chambers), as implied by the images in Fig. 4 of Fadiga and Nystul (2019). The same bias of selecting only ovarioles that had the appearance the authors expected for FSC clones appears to have been applied in the original Margolis and Spradling experiment (only a few ovarioles were selected for counting and the criteria were not stated but it is clear that the authors only considered contiguous clones occupying successive egg chambers as FSC clones), accounting for a major aspect of the artificially high contribution found in both studies.

**Figure 4.**
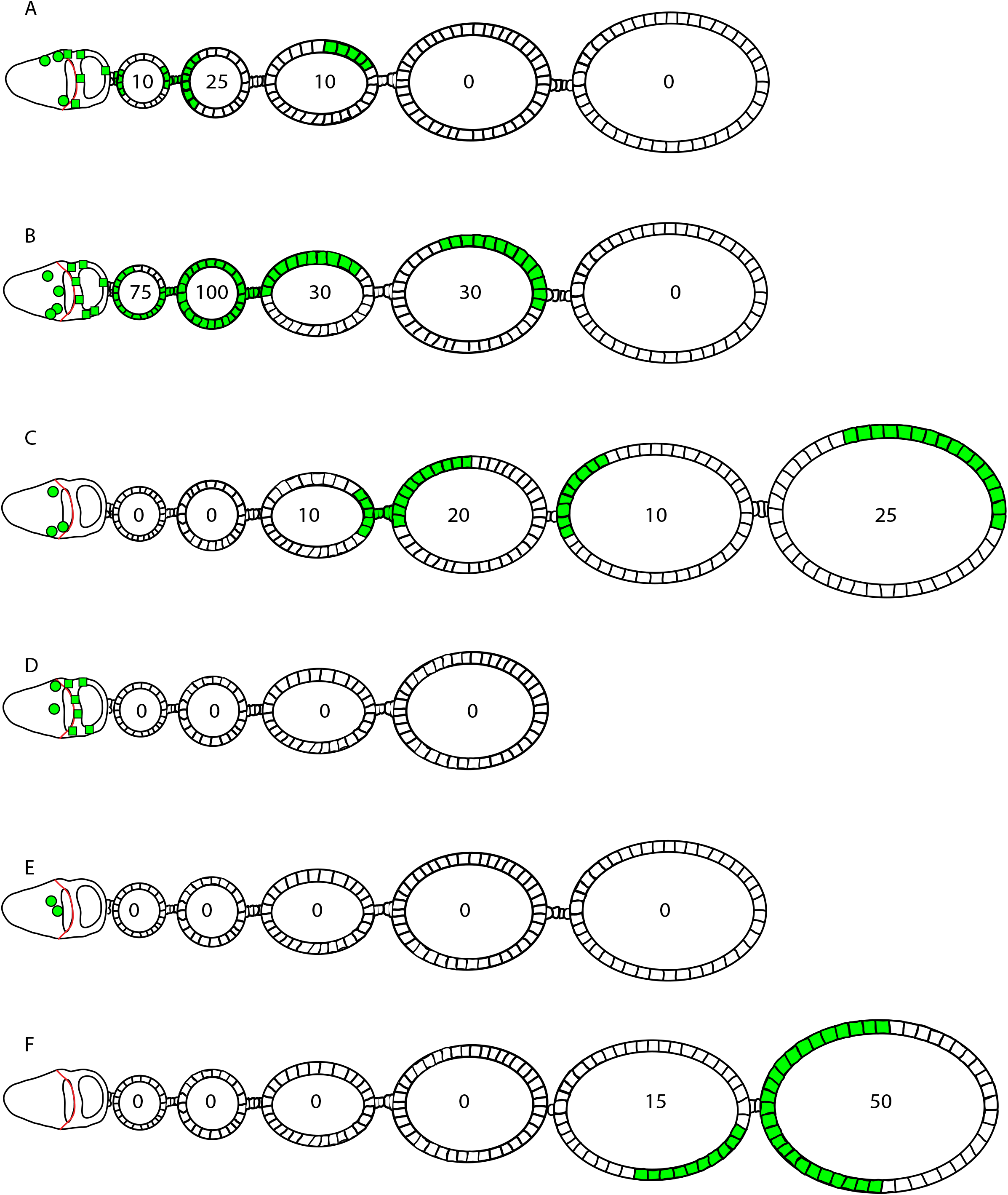
Summary of MARCM FSC lineages illustrated in Figure 3. (A-F) Diagrammatic representation of the types of FSC lineage shown in Figure 3. As explained in more detail in the legend to Figure 3, the relative frequency of the different patterns among all ovarioles examined was (A, B) 24% in sum, (C) 11%, (D) 7%, (E) 14% and (F) 9%.

The bias described above is very important to avoid with regard to FSCs and in any similar lineage analysis for any other type of adult stem cell. Even if only a single FSC is initially labeled in an ovariole, there is the opportunity for the FSC to amplify (or be lost) over time, so that the FC contribution displayed at the time of analysis represents the contribution of unknown numbers of FSCs during the intervening period (with only the final number of FSCs measured directly). This introduces a large error for FSCs (demonstrated numerically below) and would also produce a large error for any other type of stem cell maintained by population asymmetry, in which the loss and amplification of individual stem cell lineages are frequent. Importantly, the appropriate method of being inclusive rather than selective does not depend on first establishing whether stem cells are maintained by population asymmetry or single cell asymmetry, or how frequently they are lost or amplify. The correct experimental approach is not to make any assumptions about FSC behavior but simply to deduce FSC behavior by assaying, as far as possible, all ovarioles in which an FSC has been marked.

Reilein et al., (2017) performed an analysis of the proportion of the FC epithelium contributed by FSC clones for LGR multicolor analysis and MARCM analysis. Considering clones with at least one FC patch and at least one surviving FSC at 9d after heat-shock the average proportion of FCs contributed by a single clone was reported in a Supplementary Note as about 12% for RFP-negative LGR clones (chosen because it is less likely that more than one FSC was initially marked than for the more common RFP-positive genotypes) and 15% for MARCM clones. It is clear from looking at the proportion of FCs in each egg chamber for each FSC clone in both RFP-negative multicolor and MARCM clones that the average values of 12% and 15% are far lower than the 50% proposal from two FSCs of Fadiga and Nystul because (a) the vast majority of FC patches, when present, occupy less than 40% of an egg chamber (57/76= 85% for multicolor, 124/167= 74% for MARCM) and (b) a large fraction of egg chambers have no labeled FCs belonging to the FSC clone (102/178 for multicolor, 134/301 for MARCM). The average occupancy values quoted (12-15%) for FSC clones with surviving FSCs substantially overestimate the FC contribution of a single FSC for reasons that are discussed in the next paragraphs.

Had we (inappropriately) included only FSC clones that included marked FCs in the germarium as well as the first two egg chambers the average contribution from the MARCM experiment would have been calculated as 25.2%; it would have been 31.0% had we also scored only up to the third egg chamber for those clones (our best understanding of the selection and procedure used by Fadiga and Nystul), artificially inflating the result by two-fold. The fraction of FSC clones with surviving FSCs that included marked FCs in the germarium as well as the first two egg chambers in the MARCM study of Reilein et al., (2017) was 39% (24/62) at 9d, 34% (22/65) at 14d and increased to 69% (20/29) at 21d and 70% (21/30) at 28d. Correspondingly, the average number of labeled FSCs amongst all FSC clones with surviving FSCs rose from 3.3 at 9d, to 4.5 at 14d, 6.5 at 21d and 7.1 at 28d, reflecting the greater likelihood of labeled FCs being associated with consecutive cysts when there are more labeled FSCs present. These numbers also illustrate that more than 60% of the 9d clones with FCs and surviving FSCs would have been ignored using the approach of Fadiga and Nystul. Fadiga and Nystul did not disclose what fraction of FSC clones they ignored but if the distribution of clones was similar to that reported in Fig. 4B of their study (or if the FC contribution data derives from the same samples), considerably more than half were likely put aside. The above analysis of our 9d MARCM data illustrates the magnitude of the bias in the final result introduced by scoring only a subset of FSC clones (and over a limited region of the ovariole). It also makes clear that the labeled FCs in a FSC lineage with a surviving FSC will almost always be supplied during most of the time-frame scored by multiple FSCs, even if only a single FSC initially undergoes recombination. For the 9d MARCM data the contribution of about 15% of all FCs per clone was achieved by a number of FSCs that likely started as only slightly greater than one but rose to an average of 3.3 by 9d. The result is roughly consistent with an average contribution of around 6-7% per single FSC (during the period of roughly 5-8d after heat-shock when FCs are supplied to the 5-6 scored egg chambers and the average number of FSCs present is likely in excess of 2.5) that would be expected if there were 14-16 FSCs.

The estimation of the FC contribution of individual FSC clones presented in Reilein et al., (2017) based on MARCM clones 9d after heat-shock was limited to clones with a surviving FSC and proven FC-production capacity. Hence, Reilein et al., (2017) did not score clones lacking a surviving FSC or lacking any labeled FCs. Given our current understanding, in which all marked cells that are present 9d after clone induction derive from recombination in an FSC, whether those cells are just FCs, just FSCs, just ECs or any combination of these cell types, we can derive a better estimate of FSC contributions to FCs by including all ovarioles with a marked cell of any type (EC, FSC, FC alone or in any combination). That modification would reduce the measured average FC contribution to 11.3% (increasing the sample size from 62 to 117 and reducing the average number of FSCs per sample from 3.3 to 2.3). That inferred contribution of 11.3% would still be an overestimate for a single FSC because (a) we would not include ovarioles where all trace of an originally labeled FSC had been lost (a labeled FSC that became an FC in the first four days without prior duplication would not be counted) and (b) some ovarioles may initially have more than one FSC marked by recombination.

In the main text of Reilein et al., (2017) another LGR multicolor clonal analysis is reported, in which a mild single heat-shock was delivered and the range of the ovariole potentially harboring FCs from labeled FSCs of the least frequent color classes was determined and scored just 72h after heatshock, leading to a result that the average FSC lineage contributed 9.2% of FCs. The various limitations in trying to measure the FC contribution of a single FSC are least pronounced for these short-term, low frequency multicolor clones, consistent with the lowest value of 9.2% determined by this approach.

In summary, the approach of determining FSC number by scoring the contribution of single FSC clones has significant systematic limitations that will always lead to an underestimate of FSC number. Nevertheless, using more appropriate methods than those of Fadiga and Nystul (who severely biased the outcome by selecting a limited subset of FSC clones enriched for labeled FCs and counting only regions of the ovariole that were used as the selection criterion), Reilein et al., (2017) deduced that there must be an average of at least 11 (1/0.092) FC-producing FSCs per ovariole.

The key issues of FSC loss and amplification that are considered above were not discussed at all by Fadiga and Nystul; instead, the general context for consideration of which FSC clones ought to be counted is mis-represented in the prelude to presentation of experimental results. They state that “one of the original observations in the Margolis and Spradling study is that an FSC clone forms a coherent, contiguous patch of cells….. contributing to each new follicle that is formed after clone induction”. This is presented as a definitive fact or finding but the assertion was, in fact, an assumption (a generalization, extending the appearance of some clones to characterize all FSC clones) that has been shown to be incorrect in Reilein et al., (2017). No authors, including Margolis and Spradling (1995), have ever shown that all FC-containing clones present beyond the lifetime of an FC lineage (5d) have FC patches associated with every follicle; they have simply ignored the many ovarioles that do not show this pattern, and now Fadiga and Nystul (2019) confirm the description by Reilein et al., (2017) of these many other types of lineage originating in FSCs in the images and data of their Fig. 3, albeit with inappropriate names.

Fadiga and Nystul also state that “Reilein et al. posited that each FSC does not normally contribute to every follicle, so discontinuous clone patterns should be expected.” That is false. We made unbiased experimental observations and reported them, demonstrating extensive evidence from multicolor and MARCM clonal analyses for this assertion; we did not “posit” or have expectations of how clones “should” appear. It is exactly because Fadiga and Nystul made assumptions to exclude a large fraction of experimental samples, ignoring the possibility that FSCs might be lost or amplify, and we did not, that they have arrived at erroneous conclusions.

#### (iii) Location of FSCs

Fadiga and Nystul use GFP-negative clones 7d after induction to assess where FSCs are located. They report that 5/6 of all “FSC clones” have no marked cells anterior to those bordering strong Fas3 staining. In other words, they find that for 1/6 of “FSC clones” there is a putative FSC anterior to the AP location they designate as the only location of FSCs. These counter-examples are dismissed as possibly being ECs, which they call IGS cells, even for cases where there are associated labeled FCs. It is also not clear whether the samples analyzed as “FSC clones” include only clones with labeled FCs in the germarium and the next two egg chambers, as used for determination of FC contributions, or, as it should be, all clones originating in an FSC. Scoring only FSC lineages that very recently produced an FC introduces a bias against including more anterior FSCs because only the more posterior, layer 1 FSCs directly become FCs (Reilein et al., (2017) and Reilein et al., (2018)).

The biggest issue, however, is whether the authors were able to identify all GFP-negative cells in the germarium. We have found this to be extremely challenging even with extremely high quality images in our multi-color lineage analysis (because it relies on negative marking), requiring a very large investment of time, checking and re-checking permanently stored high magnification images of complete z-stacks through the germarium. It was not possible for us to accomplish that reliably and in a manner that can be checked repeatedly without generating an explicit spreadsheet record of the exact location of all marked cells in the germarium that could be FSCs. We did this for our multicolor analysis and we are therefore extremely confident of the results presented in Reilein et al. (2017), Fadiga and Nystul do not appear to have documented their findings in this manner, such that the complete catalog of labeled cells is tabulated for each sample. Additionally and importantly, we find that the objective of accurately documenting all marked cells in the germarium is far better accomplished with strong GFP-positive nuclear labeling using MARCM clones. We did exactly that in the Reilein et al., (2017) study. The results from multicolor and MARCM lineage analyses were fully concordant.

After establishing the key pre-requisite of being able to visualize and document all marked cells in potential FSC territory, it is essential to have a method for defining which marked cells are indeed FSCs. The critical criterion adopted by Reilein et al., (2017) was to examine FSC lineages in which there was only a single candidate FSC. First, regarding the radial dimension, it was found that the single candidate FSC could be found in any z-section, consistent with the radial symmetry of the germarium. Thus, FSC nuclei were always adjacent to the germarial wall but could be in any radial location. Next, FSC lineages were scored that had candidate FSCs confined to just a single AP layer (or ring). There were many examples where the only candidate FSCs were in layer 1, immediately anterior and adjacent to the border of strong Fas3 (24 FSC lineages for multicolor clones, 13 FSC lineages for 9d MARCM clones) and a similar number of examples where the only candidate FSCs were only in layer 2 or layer 3 (27 FSC lineages for multicolor clones, 16 FSC lineages for 9d MARCM clones). Also, the total number of marked FSCs (from scoring all FSC clones) in the more anterior locations (layers 2 and 3) was approximately equal to those bordering strong Fas3 staining (50%: 50% for multicolor and 46%: 54% for MARCM) and the average total number of cells labeled per germarium was 7.6 in layer 1, 5.6 in layer 2 and 2.0 in layer 3. Those results are consistent with the conclusion that there are about eight FSCs bordering strong Fas3 (layer 1) and about eight in more anterior locations (layers 2 and 3).

In Fadiga and Nystul (2019) and earlier studies the location of a FSC was not determined by looking at lineages where there was only a single candidate FSC or indeed by chronicling all candidate FSCs. Instead, it was assumed that each lineage had only a single FSC and that the FSC would be the most anterior cell in the lineage. It was never explained how only a subset of cells (roughly two out of eight) in a single AP plane could be FSCs or how those FSCs could, as claimed, always reside diametrically opposite in a mid-z-section even though there is no known asymmetry that would cause ovarioles to always lie on a slide in the same radial orientation (Nystul and Spradling, 2007; Nystul and Spradling, 2010; Reilein et al., 2017). The results of Reilein et al., (2017), showed explicitly that the single candidate FSC in multicolor clones can be at any radial location and also showed by live imaging that cells in these locations move extensively radially and exchange positions. Use of the key criterion of examining lineages with FSCs in only one candidate location, together with the volume, authentication and, most important, the acuity of the results presented by Reilein et al., (2017) showed unequivocally that FSCs mostly lie in two AP planes, far outweighing the evidence of Fadiga and Nystul claiming that FSCs lie in a single AP plane (and reporting raw data showing one in six elsewhere).

The discussion above is centered on the veracity of the conclusions of Reilein et al., (2017), showing FSCs in more than one AP plane and distributed radially along the basement membrane lining the germarial wall. The precise location of the most posterior FSCs, termed layer 1, is made very clear by images and cartoons in Reilein et al., (2017) (see Figs. 1, 2, 5 and 7 in that study). They are the cells with nuclei immediately anterior to the strong border of Fas3 expression that outlines the posterior surface of the stage 2b germline cyst. We and other labs acknowledge that there is also weaker, more anterior Fas3 expression but the anterior border of strong Fas3 expression can always be picked out in our studies (aided by the stage 2b cyst location) and has served as the key landmark from which to designate FSC locations in layers 1, 2 and 3.

Fadiga and Nystul also distinguish between strong and weak Fas3 staining and illustrate the anterior border of strong Fas3 expression with a red arrow in Fig. 5A in exactly the location we would designate. Layer 1 FSCs lie immediately anterior (to the left) of that location and, indeed, to our understanding, every study in the past has placed FSCs anterior to that Fas3 border. Remarkably, Fadiga and Nystul now indicate the cell immediately posterior (white arrow) to the strong Fas3 border as an FSC. They assert that this is generally the location of FSCs found in this study and in multiple prior studies. That assertion is not consistent with earlier descriptions from the PI. The location of FSCs has been characterized as “immediately posterior to the 2a cyst” (Nystul and Spradling, 2007) (in our analyses there can be one to three nuclei between 2a cysts and the strong Fas3 border, so cells immediately posterior to the 2a cyst could correspond to layer 1, 2 or 3 FSCs but not the more posterior location now designated by Fadiga and Nystul), by writing “not only the FSCs, but also cmcs and a variable number of additional cells near the region 2a/b border exhibit low levels of FasIII expression” (Nystul and Spradling, 2010) (low levels of “FasIII” (Fas3) in this description clearly indicated being anterior to the strong Fas3 border and Fig. 2C in that paper indicates an FSC that we would place in layer 2 or 3), by writing, “Lastly, we used other features, such as low levels of FasIII expression and a triangular-shaped nucleus to further aid in the identification of FSCs.” (Kronen et al., 2014) (again, “low level” implies being anterior to strong Fas3), and by indicating a cell anterior to the strong Fas3 border as an FSC in Fig. 1B of Kim-Yip and Nystul (2018). The new cited location of FSCs claimed by Fadiga and Nystul is very puzzling in this context, contradicting multiple earlier designations. The main point remains that FSC location must be determined by examining all FSC lineages, especially making use of lineages with FSCs in only one candidate location, and by clearly visualizing all candidate cells in an FSC lineage in the germarium relative to reproducible and reliable spatial landmarks, as was done in Reilein et al., (2017).

## 3. Discussion section in Fadiga and Nystul

In the experimental section of the paper, Fadiga and Nystul presented results from repetition of two types of experiment (and no additional, new approaches). As described above, they presented a multicolor lineage analysis that was flawed in several respects (high background clone rate, some inexplicable staining patterns and failure to induce clones at a high enough frequency to exploit the full sensitivity of the multicolor system to detect up to six distinguishable clone colors). They cite those shortcomings as “evidence” that the analyses by Reilein et al., (2017) using the same genetic system (but with clean artifact-free staining, full documentation, high clone frequencies, very low background clone frequency and no significant unexpected staining patterns) must somehow be flawed. Second, they repeated some analyses of single-color clones of the type originally conducted by Margolis and Spradling (1995) to characterize FSC clone patterns and to try to deduce FSC number by measuring FC contributions but they did so in a circular fashion, only scoring the types of clone that fit a favored hypothesis, mis-representing some of the results obtained and, additionally, using methods that do not provide clear identification and documentation of all marked cells in the germarium.

In addition to these critical deficits undermining the entirety of the experimental evidence presented, the Discussion provides further indefensible reasoning (“mis-representations” below) and fails to discuss (“major omissions” below) highly significant findings of Reilein et al., (2017) and Reilein et al., (2018) that further outline the number, location and behavior of FSCs.

### (a) Mis-representations

#### (i) Number of FSCs and FCs produced over time

Fadiga and Nystul claim that two FSCs would supply the correct number of about 900 FCs in stage 6 egg chambers and that 16 FSCs would necessarily produce too many FCs. This estimation is flawed for many reasons. Most obviously, the outcome depends critically on the rate of FSC division. Fadiga and Nystul implicitly assume that all FSCs divide every 9.6h. Not only has this not been measured explicitly for any FSCs, but it is obviously inappropriate to assume that all FSCs divide at the same rate (in a calculation for a scenario where there are 16 FSCs) when clear evidence has been presented in Reilein et al., (2017) that this is not the case (layer 1 FSCs divide more frequently than layer 2 FSCs and layer 3 FSCs rarely divide). Moreover, Reilein et al., (2018) present evidence that on average FSCs divide no more than once every 24h, far less frequently than the 9.6h Fadiga and Nystul have assumed, without explicit declaration, for their calculations based on 16 FSCs.

Second, we do not agree with the proposed time interval for FSC progeny to reach a stage 6 egg chamber and suggest that it would be more direct and accurate for evaluating the required rate of FC production to consider the shortest relevant developmental time period, namely from FSCs to a newly-budded egg chamber. Margolis and Spradling (1995) reported that egg chamber budding occurs roughly every 12h and, citing Mahowald and Kambysellis (1980), that a newly-budded egg chamber contains roughly 80 FCs; both numbers are roughly consistent with other studies. FCs first associate with a stage 2b germline cyst and most germaria have only one posterior, stage 3 cyst between that and the first budded egg chamber; occasionally there is one additional cyst. Thus, the expected time between the first “founder” FCs that associates with a stage 2b cyst and a newly budded egg chamber with about 80 FCs corresponds to about two cycles of budding, or 24h. That would allow only about three rounds of FC doubling (if FCs divide every 9.6h), so the expected number of founder FCs might, by this calculation, be ten or more. In the model proposed by Fadiga and Nystul (2019) and earlier by Nystul and Spradling (2007) there are exactly two founder FCs, one daughter from each FSC. To expand from 2 FCs to about 80 would require at least five cycles of division within the germarium, which seems implausible.

The time between founder FC association with a stage 2b cyst to development of that cyst to stage 6 has also been tested in Reilein et al., (2018) in a multicolor lineage experiment analyzed 72h after clone induction. Each recombination event produces two daughters with complementary markers retained (“twin-spots”). Recombination in a FC must always lead to both twin-spot daughter clones on the same egg chamber, whereas recombination in a FSC can lead to population of the next available germline cyst for founder FC recruitment containing only one twin-spot daughter. Reilein et al., (2018) were therefore able to determine that cells resulting from recombination in an FSC could populate the germarium and up to four egg chambers by 72h. In other words, the earliest founder FCs had progressed to the fifth egg chamber (and sometimes only the fourth) by 72h. Both the fourth and fifth egg chambers were more mature than stage 6. This evidence is in keeping with progression from founder FC to budded egg chamber in roughly 24h followed by four 12h budding cycles to progress to the fifth egg chamber, which is at stage 6 or beyond, over a total of 72h, considerably less than the 100h used by Fadiga and Nystul. In Fig. 2A of Margolis and Spradling (1995) the authors show that the contribution of FCs per clone scored in stage 10 egg chambers has saturated by day 4 after heatshock, indicating that the earliest founder FC had been labeled, and that progression from stage 6 to stage 10 takes one day, also implying a period of about 72h from founder FC to stage 6.

While the outline above seems more accurate and justifiable than the longer timeline considered by Fadiga and Nystul, both approaches include some approximations and could have significant errors. More important, therefore, is a direct experimental test of the number of founder FCs as a prelude to considering how many FSCs might produce that number of founder FCs in one cycle of egg chamber budding. We did that in Reilein et al., (2017). We found that single egg chambers in ovarioles with FSC clones that encompassed at least four colors (the time after heat-shock ensured that no clones originated in FCs) most frequently had three or more different colors (genotypes), guaranteeing at least that number of founder FCs. Several founder FCs may, however, have the same color because the palette of color diversity was limited by the number of differently marked FSC lineages (which was most commonly four and rarely as high as the maximum of six), so the observed number of colors will generally underestimate the number of founder FCs. We therefore also measured the fractional occupancy for each color. In Reilein et al., (2018) we induced FSC clones at low frequency and picked the least frequent clone colors (those lacking RFP) so that only a single founder cell would more likely be represented. In that study we found an average contribution under 19%, indicating that there are more than five founder FCs, on average, per egg chamber. In that study we also measured the average FSC doubling rate and found that it was consistent with producing the observed number of founder FCs from a population of 16 FSCs. It would obviously be extremely challenging for two FSCs to produce more than five founder FCs every 12h, requiring a much higher FSC division rate than has previously been suggested or inferred.

Although Reilein et al., (2017) and Reilein et al., (2018) present direct experimental evidence that there are at least 4-5 founder FCs per egg chamber, Fadiga and Nystul write that “multiple studies have found that transient clones are capable of covering up to half a follicle” and do not cite these contradictory papers. Even in Margolis and Spradling (1995), which is also not cited in reference to this assertion, it is stated that the average maximal clone size for transient clones in stage 10 egg chambers was 142 cells (and the total stage 10 FCs counted in that study was 651), implying 4-5 founder FCs per egg chamber. Finally, Fadiga and Nystul write that “images of very large persistent clones from many different studies… are consistent with a small number of actively dividing FSCs per germarium”. One can indeed find such images but there is very good reason to believe that these “clones” are being supported by several FSCs, not just one (see section on population asymmetry).

#### (ii) Location of FSCs

Fadiga and Spradling contest the possibility that FSCs may exist anterior to Fas3-neighboring cells by saying this “is inconsistent with many other studies that have demonstrated that somatic cells in this region have a markedly different shape, function and gene expression pattern than cells in Region 2b.” First and most important, only functional tests are relevant to defining FSCs. We reported cell shapes in Reilein et al., (2017) and noted differences in FSCs in different AP locations. We also noted differences in the magnitude of expression of some markers and in the strength of the Wnt signaling pathway. Indeed, signaling gradients likely underlie many of the differences in gene expression among FSCs. We even described differences in the behavior of FSCs in different locations, showing that posterior FSCs were more likely than anterior FSCs to become FCs in the short term. Importantly, however, we also showed that FSCs can exchange AP locations and that, over time, FSCs in all of these positions can produce FCs, as shown by the location of the sole candidate FSCs in ovarioles with FC patches in the multicolor lineage analysis of Reilein et al., (2017) (described earlier). In other words, Reilein et al., (2017) have already shown that there is heterogeneity within the FSC population but they also showed that FSCs at different AP locations, with different instantaneous properties, share the FSC-defining property of producing FCs.

#### (iii) FSC numbers and contributions

Fadiga and Nystul state, “Our study identifies several reasons for the discrepancies between the current paradigm and the Reilein et al study.” The choice of words begins the discussion by mis-representing the status quo, denying the true status and impact of Reilein et al., (2017). The conclusions of Reilein et al., (2017) and Reilein et al., (2018) were the most recent published studies directly on this topic and they therefore represented the current paradigm at the time of writing. This is not a trivial matter. Because we had to overturn the former long-standing paradigm we had to take an exceptionally thorough and multi-faceted approach, spanning many years of intensive research prior to publication. This effort was in order to uncover the best possible understanding of FSC locations and behaviors in order to benefit researchers investigating FSCs and to draw attention to general principles and new insights relevant to other types of adult stem cell. For that effort to be productive it is important that the findings are acknowledged and cited appropriately. Papers published by the Nystul laboratory specifically centered on FSC biology after the publication of Reilein et al., (2017) either did not acknowledge that work at all (Cook et al., 2017) or presented it as a minor suggested (rather than demonstrated) modification of the old paradigm (Kim-Yip and Nystul, 2018) rather than a demonstrated radical change in our understanding of FSC numbers, location and behavior; that attitude is continued here by referring to an older, overturned paradigm as the “current paradigm”. That attribution does not match the objective public domain, represented by peer-reviewed published work, and is both inappropriate and deceptive.

Fadiga and Nystul then cite four “reasons for the discrepancies”:

1. Fadiga and Nystul suggest, with no evidence, that many of the clones *we* observed in the multicolor clonal analysis of Reilein et al., (2017) resulted from epigenetic silencing because “high levels of fluorophore expression can reduce cellular fitness, so it may be that constitutive expression of three separate markers creates a selective pressure to silence one or more markers”. The LGR multicolor system only expresses two fluorescent markers (the other is *lacZ*), not three, we found as many clones lacking a non-fluorescent marker (*lacZ*) as lacking the fluorescent marker on 2L (GFP), and both the *ubi-GFP* and *ubi-RFP* transgenes are extremely widely used in ovaries and elsewhere by ourselves and others without any evidence of epigenetic silencing. More important than this unsubstantiated proposal of the cause of an alleged artifact, in our experiments the controls with no heat-shock produced virtually no background clones, confirming that the clones we observed were indeed due to *FRT*-mediated recombination induced by expression of heat-shock-inducible Flp recombinase. Fadiga and Nystul are correct in stating that “a high rate of clone induction in the absence of heat shock and the presence of clones with unexpected marker combinations are problematic for the use of clone size and frequency measurements to infer stem cell number”. There were no such issues in Reilein et al., (2017); only in the LGR experiments reported by Fadiga and Nystul.
2. Fadiga and Nystul write that the reduction in the number of FSC clones remaining in an ovariole should drift towards one over time if FSCs undergo neutral competition. We agree that is expected and it is also part of the evidence for FSC maintenance by population asymmetry. However, Fadiga and Nystul also characterize our results as showing that the average number of uniquely labeled lineages declines from “more than four at 5 dphs to two at 15 dphs, and then remain at approximately two for the subsequent 15 days”. Fig. 2j in Reilein et al., (2017) shows these average numbers declining from 2.06 at 14d to 1.89 at 20d and 1.49 at 30d, (not stabilizing at two). More important, it is essential to look at the results for each ovariole, not just averages. The results for every ovariole are supplied in the Supplementary Note of Reilein et al., (2017) and show that the percentage of ovarioles with only one color of FSC clone increases from 17% at 14d to 30% at 20d and 56% at 30d. This is exactly what is expected from population asymmetry and neutral drift (and directly contradicts the portrayal of our findings by Fadiga and Nystul). Fadiga and Nystul also write that they “found no evidence for the claim that individual FSC clones can be non-contiguous and contribute only sporadically to follicles..” In fact, they highlight such a clone in Fig. 1F and show another two examples (the clone outlined in white in Fig. 1B does not populate at least two egg chambers, the clones highlighted in white and in magenta each miss one egg chamber [and only half of the ovariole is typically shown]) in their multicolor analysis. They also describe and show four categories of such clones in their GFP-negative clonal analyses (which they term as Transients, FSC loss, Small and Discontinuous but all result from recombination in an FSC, as explained earlier), representing at least 40% of all ovarioles. Clearly the statement of “found no evidence” does not accurately represent their results. Fadiga and Nystul acknowledge that Reilein et al., (2017) also deduced there were many more than two FSCs by measuring the average contribution of a single FSC. They write, however, that the representation of lineages derived from FSCs in different locations may be unequal because the frequency of marking by FRT-mediated mitotic recombination depends on proliferation rate. We agree that is conceivable. From our understanding of FSC behavior, the net effect would be to reduce the representation of the more anterior, slower-dividing FSCs among initially labeled FSCs. Those FSCs must move posteriorly before they can become FCs and therefore are likely to contribute the fewest FCs. So, this consideration may lead to an over-estimation of the FC contribution of an average FSC and hence an under-estimation of FSC number. It cannot account for our estimation of FSC number from this approach being too large. The issue is also bound to have a only a very small impact because FSCs are continually exchanging AP locations. In the Fadiga and Nystul postulate of two FSCs dividing at equal rates the consideration they raise would have no bearing on the representation of different FSCs among marked clones. Fadiga and Nystul are therefore asking readers to consider the impact of differing rates of FSC proliferation and are, at the same time, saying that there are no such differences.
3. Fadiga and Nystul repeat the incorrect assertion that Reilein et al., (2017) assumed that *FRT* recombination occurs at both sites in 100% of mitotic cells. This assertion is clearly wrong both because we made no such assumption (discussed earlier) and because the outcome of that scenario would be the absence of clones with both *lacZ* and GFP (because recombination leads to loss of one marker or the other) and reduction of the total possible number of different clone types to four (neither of which were close to what we observed). The thinking behind this assertion is clearly confused. In addition, “By assuming that the recombined phenotypes (marker recombinations) should be more common than they actually are…” is a mis-representation; we scored the actual, observed recombined phenotypes and then used a statistical model.
4. Fadiga and Nystul measure clone size using single-color lineage analysis, report a similar result to the original Margolis and Spradling (1995) study and imply that our measurements of the same parameter must therefore be erroneous (for unspecified reasons). We described earlier how Fadiga and Nystul made a fundamental error of selecting a subset of ovarioles and a limited region within those ovarioles to score rather than including all ovarioles that initially included a labeled FSC and scoring as many egg chambers as possible in each such ovariole, as we did for both multicolor and MARCM analysis. The bias from selecting only clones meeting expectations of continuous large FC contributions greatly skews the results; the underlying reason is that most such clones derive from several FSCs, not just one, even if in many cases a single FSC initially undergoes recombination (FSCs in most surviving lineages amplify over time as a fundamental feature of neutral competition).

### (b) Major omissions

Fadiga and Nystul focus on the question of how many FSCs are in a germarium. They do not explicitly discuss the intertwined additional major insights from Reilein et al., (2017) that (i) FSCs are maintained by population asymmetry and that (ii) FSCs produce ECs as well as FCs.

#### (i) Population asymmetry and neutral drift

The multicolor lineage analysis of Reilein et al., (2017), examining the number of distinguishable surviving FSC lineages at different times after induction showed a rapid reduction of FSC lineages over time, contrasting with earlier proposals that the average half-life of each FSC was over two weeks. That observation, coupled with the demonstrated increasing number of FSCs representing each surviving lineage over time, and the observed stochastic distribution of the number of FSCs per lineage at any one time over a number of ovarioles, are characteristic of population asymmetry and neutral competition among FSCs (Jones, 2010; Blanpain and Simons, 2013). The rapid loss of marked FSC lineages, especially shortly after induction when many lineages are still represented by only a single FSC, was further confirmed in Reilein et al., (2018) by directly measuring the loss of 25 out of 49 FSC lineages over 3d. The reason that multiple earlier studies greatly overestimated FSC stability is because they invariably looked at loss of FSC lineages from a starting point of 7d or more after clone induction. By the starting point of those experiments many FSC clones have been lost (hence, those ovarioles are not scored) and the number of FSCs in the surviving lineages has increased, so that subsequent extinction of a lineage requires the loss of multiple marked FSCs, not just one.

Fadiga and Nystul do not engage with the possibility and implications of population asymmetry for FSCs. They write: “Indeed, if FSC clones were typically discontinuous, it would be very difficult to distinguish between persistent FSC clones and transient clones …”. This reveals a belief that FSC clones must be persistent as a defining property; that is an incorrect assumption. The definition of FSCs is not a matter of subjective preference, to be altered in order to disregard some FSCs or to suggest that there is no objective way to define FSCs. FSC clones must be distinguished from FC clones purely on the basis of timing, without assuming what such lineages should look like. The inappropriate assumption that an FSC clone is defined by continuous, extensive FC production, which turns out to be far from correct, is evident in earlier work: “..restricted our analysis to ovarioles with mature FSC clones, defined as those that originate at the region 2a/2b border and include roughly half of the follicle cells in the germarium.” (Kronen et al., 2014) and “FSC clones can be identified as large contiguous groups of marked cells that extend from the region 2a/2b border through the germarium and into the budded follicles” (Cook et al., 2017).

Fadiga and Nystul also write, “these transient clones may have been misinterpreted as evidence for persistently labeled FSC lineages”. Not so. The referenced clones are indeed, by the key functional definition, FSC lineages but they are not necessarily persistent; their persistence is variable and stochastic. Fadiga and Nystul “favor the alternative hypothesis that the clonal diversity at early time points is due to the presence of transient clones that had not yet moved out of the tissue”. The term “transient clones” is extremely mis-leading. It refers to a vaguely defined phenotype (no specific time limits on the transience) but the only relevant question is in which type of cell the clone arose (in which cell recombination took place). An FC (sometimes called a pre-follicle cell when still in the germarium) is by definition a cell that associates with a germline cyst and must therefore move out of the germarium with the germline cyst. All such cells and their FC progeny will have moved out of the ovariole before we score clones in all of our analyses (only 5d is required for this but we use later time points). All marked FC patches observed more than 5d after clone induction therefore result from recombination in a cell that is upstream of an FC. All such cells were originally defined as stem cells in Margolis and Spradling (1995) and those upstream cells have been called stem cells by all investigators over the last 25 years. The designation also accords with the definition of all types of adult stem cells as the cells that are collectively responsible for maintaining production of one or more specific cell types (FCs in this case) throughout adult life (Clevers and Watt, 2018; Post and Clevers, 2019). This designation does not preclude subsequent discoveries of two or more types of FSC (considered further below) or additional cells that may become FSCs under specific conditions of environmental or genetic manipulations. However, it does preclude calling some fraction of functionally defined FSCs that operate during normal physiological conditions by some other name.

The historical evolution of ideas about FSCs has many similarities to the paradigm of mouse intestinal stem cells. Early lineage studies focused on long time periods, by when each developmental unit (ovariole or crypt) was populated by one or a very small number of lineages. Subsequent experiments included the key features of looking at much earlier elapsed times, distinction between lineages in the same developmental unit by using multiple colors, comprehensive analysis of all cells and all samples, using the core definition of a stem cell, and putting aside earlier bias or misconceptions (Jones, 2010; Clevers and Watt, 2018; Post and Clevers, 2019). In both cases these advances revealed that there were multiple stem cells, which underwent rapid loss or expansion in a stochastic manner, characterized as population asymmetry. In neither case is it appropriate to maintain the discredited older concept of a small number of stable stem cells or to dismiss the observations underlying these advances by inventing another name to refer, arbitrarily, to some or most of the stem cell population.

##### Further considerations of population asymmetry

It is evident that the term “population asymmetry” is not uniformly understood or always used to convey the same idea with reference to adult stem cells. The wider context of this issue is that adult stem cell function can only adequately be defined in terms of a group of cells (which maintain production of one or more derivative cell types throughout adult life) (Clevers and Watt, 2018; Post and Clevers, 2019). Experimentally, however, it is generally necessary to define which cells are stem cells (how many and in what locations) by lineage studies. Lineage studies report what happened to individual cells over time. This can lead to incorrect conclusions if not considered carefully. For example, if two cells, which may be equivalent in their initial location, properties and potential, are traced, one lineage disappears quickly and the other is long-lived it may be tempting to conclude that only the latter is a stem cell lineage because only the latter appears to be contributing to production of derivatives (potentially) throughout adult life. The appropriate treatment of lineage analyses is explored below.

It is useful to consider a thought experiment in which a developmental unit (isolated section of tissue) that is maintained by stem cells contains an array of cells, labeled, for example, 1-1 through 1-8 and 2-1 through 2-6, to report their locations in two dimensions (the analogous labels for FSCs would roughly correspond to eight possible radial locations of layer 1 FSCs and 6 radial locations of layer 2 FSCs). If we imagine being able to follow the behavior of all of these cells live over an extended period of time we might find, for example, that at the end of a given, appropriate time period (discussed further below) that lineages initiated by cells 1-3, 1-4,1-5, 2-1 and 2-3 were still present but lineages from the other nine cells were lost from the developmental unit (such as an ovariole). This should not be interpreted as evidence that only some of the 14 cells (specifically those in the named five locations) are stem cells. Instead, it is important to assay many developmental units in the same experiment and to pool the results. We may find that the lineages that survive in other developmental units derive from different locations and that the number of surviving lineages is not always the same. If, in the collective data, we find that all or a specific subset of cell positions give rise to surviving lineages at similar frequencies we should conclude that a stem cell can be at each of those locations and that cells in each location have a similar functional contribution on average. We would therefore designate all of those cells as stem cells in the knowledge that each has the potential to contribute derivatives long-term but that only a subset (that cannot be forecast) will do so.

Lineage analyses of the type performed in Reilein et al., (2017) roughly mirror the thought experiment of live tracing, but they document only the final location of cells. However, when there is a surviving FSC lineage (defined by the time that has passed) with only a single cell at a plausible stem cell location it is reasonable to infer that a stem cell can reside at that location (the inference will not always be correct because a stem cell may sometimes no longer be present). By recording those locations for many lineages over many ovarioles it was found that stem cell locations were found at similar frequencies in layer 1 and in layer 2, and for all layer 1 and layer 2 radial positions, leading to the deductions that all of these 14 locations harbor FSCs. The observations that each individual ovariole in the experiment showed a continuing contribution from only a few of these FSC locations, that some lineages without surviving FSCs still had traces of the lineage (as labeled FCs) and others lacked any representation at the time of analysis, should not be interpreted as showing that only a subset of the 14 cells are stem cells or that the remainder can be assigned identities as transient progenitors.

In both the thought experiment and the lineage tracing experiments the period of time between the start and end of the experiment is very important. In some tissues it may be hard to be certain what period is sufficient to exceed the lifetime of a derivative cell and therefore ensure that all derivative cells present at the end of the experiment were produced from a cell that was an upstream stem cell at the start of the experiment. The ovariole presents a very favorable case because the derivative FCs are defined by their association with germline cysts and inevitable passage through the ovariole with a well-established timeline. That allows an experiment to be designed such that all derivatives examined at the time of analysis originated from a stem cell at the time a lineage was initiated. In order to be absolutely sure of this, the time interval used in the Reilein et al., (2017) studies was at least 9d, at least 4d more than strictly necessary. The consequence of this choice is that all of the lineages reported can certainly be assigned a FSC origin and there is no question of giving those upstream cells a name other than FSCs. In ovarioles all FC-producing transient clones are clones that initiate in an FC and they cannot, by definition of an FC, persist for longer periods than 5d.

While the observations discussed above showed a population of FSCs with a key shared, defining stem cell behavior, might there be sub-divisions within the FSC population? Reilein et al (2017) in fact went further, finding that the immediate characteristics and behavior of FSCs was different depending on their precise AP location (most crucially, direct production of ECs by anterior FSCs and direct production of FCs by posterior FSCs). However, they also found that FSCs can exchange AP locations and that a lineage derived from a single FSC can, over time, include behaviors characteristic of each AP location (production of both ECs and FCs). These observations, discussed in greater depth below, provide clear evidence that, even though there is clear heterogeneity within the FSC population, all of the FSCs defined by Reilein et al., (2017) behave as a single community and must all, accordingly, be given the same name of FSCs.

#### (ii) FSCs produce ECs as well as FCs

Lineage studies that permit thorough scoring of all marked cells in the germarium have shown that adult ECs (termed IGS cells by Fadiga and Nystul), as well as FCs can be labeled by *FRT*-mediated recombination events in adults (Decotto and Spradling, 2005; Kirilly et al., 2011; Reilein et al., 2017). Decotto and Spradling proposed that the labeled cells in the anterior half of the germarium derived from an Escort Stem Cell located adjacent to Germline Stem Cells at the extreme anterior of the germarium. That proposal was later rejected by Morris and Spradling, and by Kirilly et al.; it was always at odds with the common finding of no measurable somatic cell proliferation in Region 1 (the anterior third of the germarium) (Kirilly et al., 2011; Morris and Spradling, 2011; Reilein et al., 2017). Kirilly et al. instead suggested that the source of labeled ECs was the subset of ECs located in Region 2a close to Region 2b (Kirilly et al., 2011). That region, encompassing the maximal anterior extent of proliferating somatic cells, corresponds to our designation of layer 2 and 3 FSCs (and some of layer 1) (Reilein et al., 2017).

The key issue was therefore whether these EC-producing cells were restricted to EC production or could also produce FCs. Kirilly et al. and other investigators had postulated that EC lineages and FC-containing lineages were distinct, resulting from recombination in different cells even when both types of clone were found in the same ovariole (Decotto and Spradling, 2005; Kirilly et al., 2011). Reilein et al (2017) showed that the proportion of ovarioles with a marked EC increased over time, so that a cell other than a labeled EC must be the source. Reilein et al., (2017) also found that the proportion of FC-containing clones that also contained one or more marked EC rose over time such that almost all FC-containing ovarioles also included marked ECs by 22 days after clone induction. In other words, virtually all FSCs that produce FCs and survive will eventually produce ECs. The clear implication that a single cell can give rise to a lineage containing both FCs and ECs was confirmed by labeling FSCs at very low frequency to ensure that most clones of a specific color combination arose from a single cell (Reilein et al., 2017). In these experiments the products of multiple single cell lineages were captured 3d after induction. Many of these lineages included both ECs and FCs, and there were also examples of FC-lineages and EC-lineages with complementary “twin-spot” colors indicating a shared parent. Thus, a single FSC can produce both FCs and ECs.

Importantly, Reilein et al., (2017) also correlated recent FC production with FSC location and directly observed movement of FSCs into EC territory by live imaging to conclude that ECs derive directly from anterior FSCs, while FCs derive directly, entirely or predominantly, from posterior, layer 1 FSCs. These observations also explain why clones containing only FCs or only ECs can readily be seen at early times after clone induction (when the parent FSC and its derivatives may be confined to a single layer). Later, most FSC lineages that survive have amplified and occupy more than one layer. Reilein et al., (2017) induced multicolor clones at low frequency and used additional criteria to be confident that almost all derived from single cells. Among those lineages containing two or more FSCs when examined 3d after clone induction, two thirds (16/24) spanned more than one layer, showing directly that one or more daughters of the original FSC must have moved, or been projected, from one layer to another. The movement of FSCs between layers appears to occur in both directions. For the five lineages that had produced an EC and still retained one or more FSCs there was an FSC in layer 1 in four cases. For lineages that had produced only FCs, suggesting the original FSC was in layer 1, and still included one or more FSCs, there was an anterior (layer 2 or 3) FSC in eight of nine cases. In five of these cases the lineage included FCs in the first egg chamber available to marked FSCs, making it almost certain that the original FSC was in layer 1. Similarly, of the nine lineages that included both an EC and FCs, seven had contributed FCs to the first available egg chamber, indicating the initial presence of a FSC in layer 1, early production of FCs and production of an EC only later. Thus, over time, an FSC lineage that initially produced only FCs could also produce ECs and vice versa. These insights clearly reveal that the entire FSC community, including FSCs at different AP locations, must be considered as one population with the potential to produce both FCs and ECs, even though at any one time the more posterior FSCs divide faster on average, experience different magnitudes of signaling from important external signals such as Wnt, Hedgehog and JAK-STAT ligands, have correspondingly different magnitudes of marker gene expression and have different instantaneous behavior options available.

Fadiga and Nystul (2019) completely ignore these results and conclusions, instead citing only the studies of Kirilly et al. to assert that labeled cells in FSC lineages that they find in more anterior positions than their preferred designated FSC location may be IGS cells unrelated by lineage to the FSC clone in the same ovariole.

#### (ii) Division-independent differentiation

Reilein et al., (2018) used the multicolor lineage system (and also mutations that affect FSC division rate) to determine whether the production of an FC is coupled temporally or functionally to FSC division. They found that FSCs divide to produce two FSCs and that an FSC can become an FC at any time relative to its last division. These results are further confirmation that FSCs are maintained by population asymmetry but also reveal more mechanistic detail about the type of population asymmetry. Most important, theoretical analysis was used to show that in all systems where stem cells are maintained by division-independent differentiation that neutral competition amongst stem cells depends on equal rates of proliferation. Any stem cell with a heritably altered rate of proliferation will compete less will if it divides more slowly and it will compete better if it divides faster than other FSCs. That prediction was tested experimentally (and found to be correct) for FSCs by measuring the loss and amplification of FSC lineages with a variety of genetic alterations that decrease or increase division rate. It is important to note that the dependence of FSC longevity and amplification on proliferation rate is only expected for division-independent differentiation, and not for other potential modes of population asymmetry and not if stem cells act independently (single cell asymmetry) as originally proposed for FSCs by Margolis and Spradling (1995), and supported by Fadiga and Nystul (2019). Thus, the discovery that FSCs are maintained by population asymmetry in Reilein et al., (2017) was verified and explored further in Reilein et al., (2018), and it was shown to provide an explanation for the fact that one of the major parameters controlling representation and longevity of an FSC lineage is the rate of proliferation (Wang and Kalderon, 2009; Wang et al., 2012; Huang and Kalderon, 2014). That experimental finding would require an entirely different (and unknown) explanation if each FSC lineage were instead long-lived and not subject to population asymmetry, as the old, incorrect FSC paradigm had stated.

These additional insights of Reilein et al., (2017) and Reilein et al., (2018) are directly relevant to, and fully support the conclusions of Reilein et al., (2017) regarding the issues of FSC number, location, longevity and output that Fadiga and Nystul are questioning. Instead of acknowledging all of these additional studies, which clearly explain (with evidence) how FSCs produce both ECs and FCs, and how FSCs are stochastically maintained by population asymmetry, Fadiga and Nystul suggest that they have discovered some “flexible, stochastic FSC behaviors” that will be interesting to explore in the future together with the possibility of finding some conditions under which “cells can convert from the IGS cell identity into a follicle cell identity or vice versa”. Thus, not only do Fadiga and Nystul ignore extensive further evidence relevant to the issue of FSC number and productivity but they also use the same device in an attempt to re-write history concerning other key FSC behaviors already demonstrated in Reilein et al., (2017) and Reilein et al., (2018).

## Conclusions

Three major conclusions in Reilein et al., (2017) drastically revised understanding of the basic arrangement and behavior of FSCs in the Drosophila ovary: (i) there are 14-16, rather than two or three FSCs per germarium, (ii) these FSCs reside in all radial locations within a narrow AP domain of each germarium, and (iii) they are maintained by population asymmetry with frequent loss and amplification of individual lineages rather than repeated asymmetric division outcomes for each FSC. We have described in detail the experimental evidence and logic that underlies these important conclusions and why the counter-claims of Fadiga and Nystul (2019) are comprehensively flawed and do not detract from these conclusions in any way. We also described how further foundational findings in Reilein et al., (2017) and Reilein et al., (2018) elaborate and build on the first three conclusions, showing that (i) FSCs, defined by the production of proliferative Follicle Cells (FCs), also produce non-proliferative Escort Cells (ECs), which support early germline differentiation in the anterior half of the germarium, (ii) posteriorly located FSCs directly become FCs, while anterior FSCs produce ECs but FSCs also change AP locations relatively frequently so that a single FSC lineage can produce both FCs and ECs, (iii) the strength of Wnt signaling, which declines from anterior to posterior across the FSC domain, strongly influences FSC location and the propensity to produce ECs or FCs, and (iv) that FSCs undergo division-independent differentiation, with the important consequence that a stem cell with a heritably altered rate of proliferation competes less will if it divides more slowly and competes better if it divides faster than other FSCs.

The most fundamental flaw in the Fadiga and Nystul (2019) study is a failure to define an FSC and an FSC lineage appropriately, and the consequent inadequacy of the experimental steps necessary to detect and characterize the quantitative properties of all FSCs and their derivatives. Specifically, in repeating a multicolor lineage analysis designed by Reilein et al., (2017), Fadiga and Nystul did not achieve a high enough frequency of recombinant genotypes or score a sufficient segment of each ovariole to count the number of distinguishable FSC lineages present as sensitively as in Reilein et al., (2017), resulting in a significant under-count. Fadiga and Nystul also reported defects in their multicolor lineage analyses to allege, without any direct evidence, that the genetic system itself might be flawed. However, none of these defects were present in the studies of Reilein et al., (2017) or in a recent repetition.

In the remaining experimental portion, Fadiga and Nystul use single color lineage analysis but repeat a key flaw of several studies prior to Reilein et al., (2017) by sampling only the fraction of FSC lineages conforming to their preferred hypotheses. By inappropriately excluding a large proportion of FSC lineages, the authors greatly overestimate the average Follicle Cell contributions of each FSC lineage and incorrectly attribute the products of each lineage to a single FSC. Finally, Fadiga and Nystul sought to establish the location of FSCs through single-color negative marking. That approach did not permit the thorough scoring and recording of all labeled candidate FSCs in every lineage and it did not use the crucial criterion of scoring lineages with only a single candidate FSC location. Those objectives were fulfilled in Reilein et al., (2017) with fully concordant results from multicolor lineage analysis and positive single-color lineage analysis, which provides the clearest images of all marked cells in the germarium. The results showed unequivocally that FSCs reside principally in two anterior-posterior planes and at each radial location within those planes, fully consistent with the estimate of about 14-16 FSCs. Fadiga and Nystul additionally mis-state or mis-represent many of the key concepts, results and conclusions of Reilein et al., (2017) that they criticize, while ignoring a number of other findings from that paper and Reilein et al., (2018) that are directly relevant to the topic under consideration.

## Notes

### Competing Interest Statement

The authors have declared no competing interest.

