## Supplemental Figures for "Follicle Stem Cells (FSCs) in the Drosophila ovary; a critique of published studies defining the number, location and behavior of FSCs"

### Supplementary Figure Legends

**Supplementary Figure 1. Additional MARCM FSC lineages with marked FSCs and FCs.**
(A-J) A selection of MARCM clones from a single experiment examining ovarioles 9d after clone induction. Panels on the left include the germarium and early egg chambers, while panels on the right (and center) show the remainder of the same ovarioles, with numbered egg chambers to make clear overlapping regions. GFP-labeled cells (green) in each lineage are shown together with DAPI staining of all nuclei (blue) and Fas3 antibody staining (red) in select panels. Each image includes only a limited set of z-section, so not all GFP-positive cells are visible. FSCs are indicated by colored arrows in layer 1 (white), layer 2 (blue) and layer 3 (yellow). The estimated proportion of the FC epithelium of each egg chamber that has GFP label is estimated by the indicated percentage values (calculated by looking through all z-stacks). (A-F) Additional examples of ovarioles with marked FSCs and FCs in both the germarium and one or more egg chambers. Included are examples where FSCs are (A) in only the posterior layer (layer 1) or (E) only in anterior layers 2 and 3. (G, H) An ovariole with marked FSCs and FCs in one or more egg chamber but not in the germarium. (I, J) An ovariole with marked FSC but no FCs. Marked cells anterior to the FC (left) are ECs.

**Supplementary Figure 2. Additional MARCM FSC lineages with marked FCs but no FSC.**
(A-D) A selection of MARCM clones from a single experiment examining ovarioles 9d after clone induction. GFP-labeled cells (green) in each lineage are shown together with DAPI staining of all nuclei (blue) and Fas3 antibody staining (red). Each image includes only a limited set of z-section, so not all GFP-positive cells are visible. A white arrow indicates the most anterior location of GFP-positive cells (other than the ECs in the anterior of the germarium in (B) and (C)). Amongst all such ovarioles, the most anterior GFP-positive cells were found with similar frequency in association with the posterior germarial cysts and each of the subsequent egg chambers, suggesting that (A-D) capture loss of the last FSC (to become an FC) (A) in the last budding cycle, (B) two cycles earlier, (C) three cycles earlier and (D) four cycles earlier.

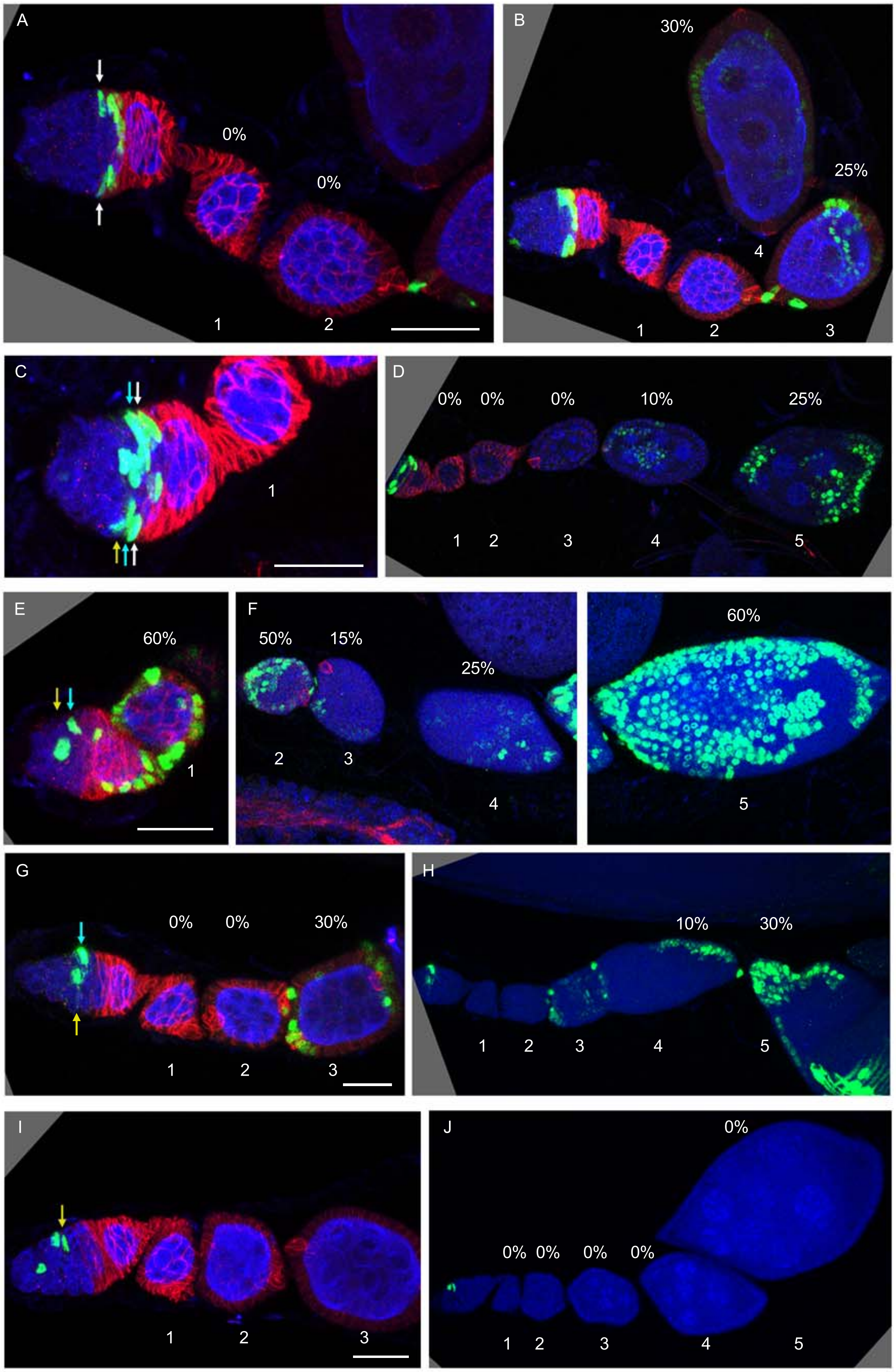


Suupplementary Figure 1

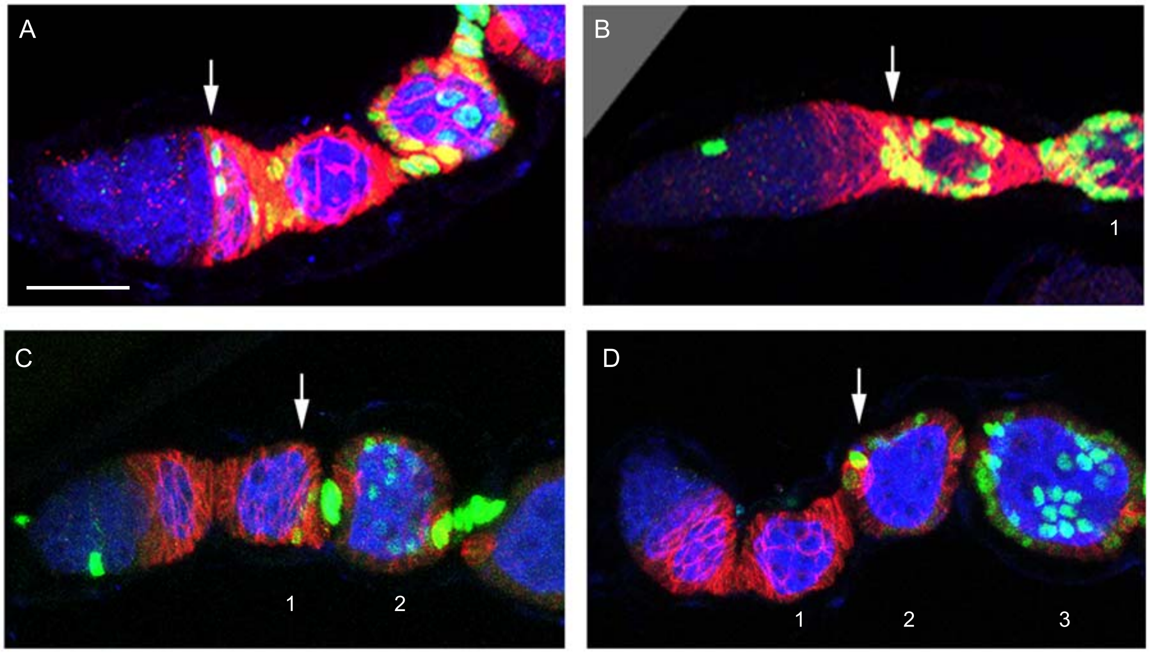

Supplementary Figure 2
